# AllTheBacteria: a community resource empowers biology and discovers novel peptide antibiotics

**DOI:** 10.1101/2024.03.08.584059

**Authors:** Martin Hunt, Marcelo D. T. Torres, Nabil-Fareed Alikhan, Daniel Anderson, Maria Luiza Andreani, Josefin Blom, George Bouras, Fiona S.L. Brinkman, Laura M. Carroll, Matthew A. Croxen, R. Andres Floto, Michael B. Hall, Jane Hawkey, Samuel T. Horsfield, Baofeng Jia, Jake A Lacey, Hyun-Su Lee, Leandro Lima, Neil MacAlasdair, Sudaraka Mallawaarachchi, William Matlock, Ahmed M. Moustafa, Robert Petit, Vignesh Ramnath, Vishnu Raghuram, Matthew J. Russell, Theo Sanderson, Timo Saratto, Oliver Schwengers, Torsten Seemann, Liam P. Shaw, Wei Shen, Nicholas Thomson, Gerry Tonkin-Hill, Jackie Toussaint, Thanh Le Viet, Johanna von Wachsmann, Fangping Wan, Aaron Weimann, Rachel M. Wheatley, Maciej Wiatrak, Ouli Xie, Cesar de la Fuente-Nunez, John A. Lees, Zamin Iqbal

**Author notes:** Corresponding authors: Cesar de la Fuente-Nunez; John A. Lees; Zamin Iqbal. Contributed equally (Martin Hunt, Marcelo D. T. Torres). Contributed equally (Cesar de la Fuente-Nunez, John A. Lees, Zamin Iqbal).

## Abstract

Public microbial genomes encode an immense record of biological diversity, evolution and molecular function, but much of this information remains difficult to reuse because raw sequencing data are not uniformly assembled, quality controlled, annotated or searchable at scale. Here we present AllTheBacteria, an open, community-built resource that transforms public bacterial short-read whole-genome sequencing reads into a uniformly processed discovery platform. The current analysed release contains 2,440,377 high-quality bacterial and archaeal genomes from 11,273 species, together with standardized taxonomic assignments, genome annotations, antimicrobial resistance calls, antiphage-defence annotations, protein structure predictions and AI-ready sequence tables. We show that this infrastructure enables applications that would otherwise be impractical, from global sequence search and outbreak contextualization to pangenome method development, antimicrobial resistance reservoir mapping and antiphage-defence ecology. As a stringent experimental demonstration, we mined 3,919,096 encrypted peptide fragments from AllTheBacteria proteomes using our deep learning model APEX 1.1, identifying 1,867 candidates with predicted antimicrobial activity. We synthesized 24 representative peptides and tested them against 20 clinically relevant bacterial strains, including antibiotic-resistant pathogens. Multiple peptides showed low-micromolar activity, membrane-responsive conformational transitions and selective envelope perturbation. A lead molecule, ATB20, reduced *Acinetobacter baumannii* burden in a murine skin abscess model with efficacy comparable to polymyxin B and no overt toxicity. Together, these results establish AllTheBacteria as both a foundational community resource for microbiology and a renewable engine for AI-guided antimicrobial discovery.

---

The rapid expansion of bacterial and archaeal sequencing has created an unprecedented opportunity to study evolution, ecology and function directly from genomes. Bacteria and archaea shape every ecosystem on Earth and include both beneficial organisms and pathogens responsible for major infectious disease burdens. The public record of bacterial genomes should, in principle, reveal reservoirs of antimicrobial resistance, antiviral immunity, biosynthetic potential, metabolic innovation and molecules that may be repurposed for therapy. In practice, this promise has been constrained less by sequencing than by reuse: the information needed for discovery is distributed across raw archives, inconsistent metadata, heterogeneous assemblies and non-standardized downstream analyses.

Large, harmonised genomic resources can expand the scale of what the research community can achieve. In human genetics, ExAC^1^ and gnomAD^2^ aggregated hundreds of thousands of genomes into standardised datasets that enabled systematic measurement of genetic constraint, improved variant interpretation and accelerated biological and therapeutic discovery^3^. Microbiology needs an analogous resource, but the problem is more complex: bacterial genomes vary not only by point mutation and vertical inheritance, but also by rapid gene gain, gene loss and horizontal transfer through interaction with mobile elements. As a result, a single species can harbour huge genetic diversity, making comprehensive sampling and standardized processing essential for understanding bacterial and archaeal biology at planetary scale.

International Nucleotide Sequence Database Collaboration (INSDC) databases archive raw sequencing reads from bacterial submissions, which are a critical fundamental resource, but they but do not consistently convert them into harmonised assemblies with standardised quality metrics, taxonomic assignment, annotations or searchable derived data products. As a result, many basic operations (e.g. searching millions of genomes for a gene, extracting all close relatives of a query genome) and focussed studies (e.g. comparing defence-system repertoires or training AI models on microbial genome context) remain limited to groups with substantial computational capacity and bespoke pipelines. Analyses can be influenced by differences in gene annotation^4^. Existing assembly efforts have made important progress^5,6^ but have not yet provided a fully open, community-extensible infrastructure designed simultaneously for microbiology, evolutionary genomics, public-health surveillance, structural biology and AI.

We therefore created AllTheBacteria, a community-governed atlas of public bacterial and archaeal isolate genomes that is designed not merely as a static database, but as a living discovery system. AllTheBacteria centrally processes public sequence data using transparent workflows, standardizes quality control and taxonomic assignment, exposes data products in formats usable by different research communities and supports continual extension by domain experts. Here we describe the resource, demonstrate its value across diverse biological questions and use it to show bacterial proteomes contain a large reservoir of encrypted antimicrobial peptides that can be recovered by deep learning and validated experimentally. This combination of resource building, algorithmic infrastructure and wet-lab validation is the central advance of the study.

## A community-built atlas of bacterial and archaeal genomes

AllTheBacteria ingests short-read isolate sequencing data labelled as bacteria or archaea in INSDC and converts them into high-quality, analysis-ready genomes. The current release analysed 2,440,377 genomes from 11,273 species. Although the collection is naturally biased toward organisms most often sequenced in research and surveillance – especially human pathogens – it captures a broad spectrum of public bacterial and archaeal diversity and is composed overwhelmingly of natural isolates rather than laboratory experiments (Fig. 1a,b). This breadth allows the resource to support both focused species-level studies and large-scale analyses across the tree of life.

**Figure 1.**
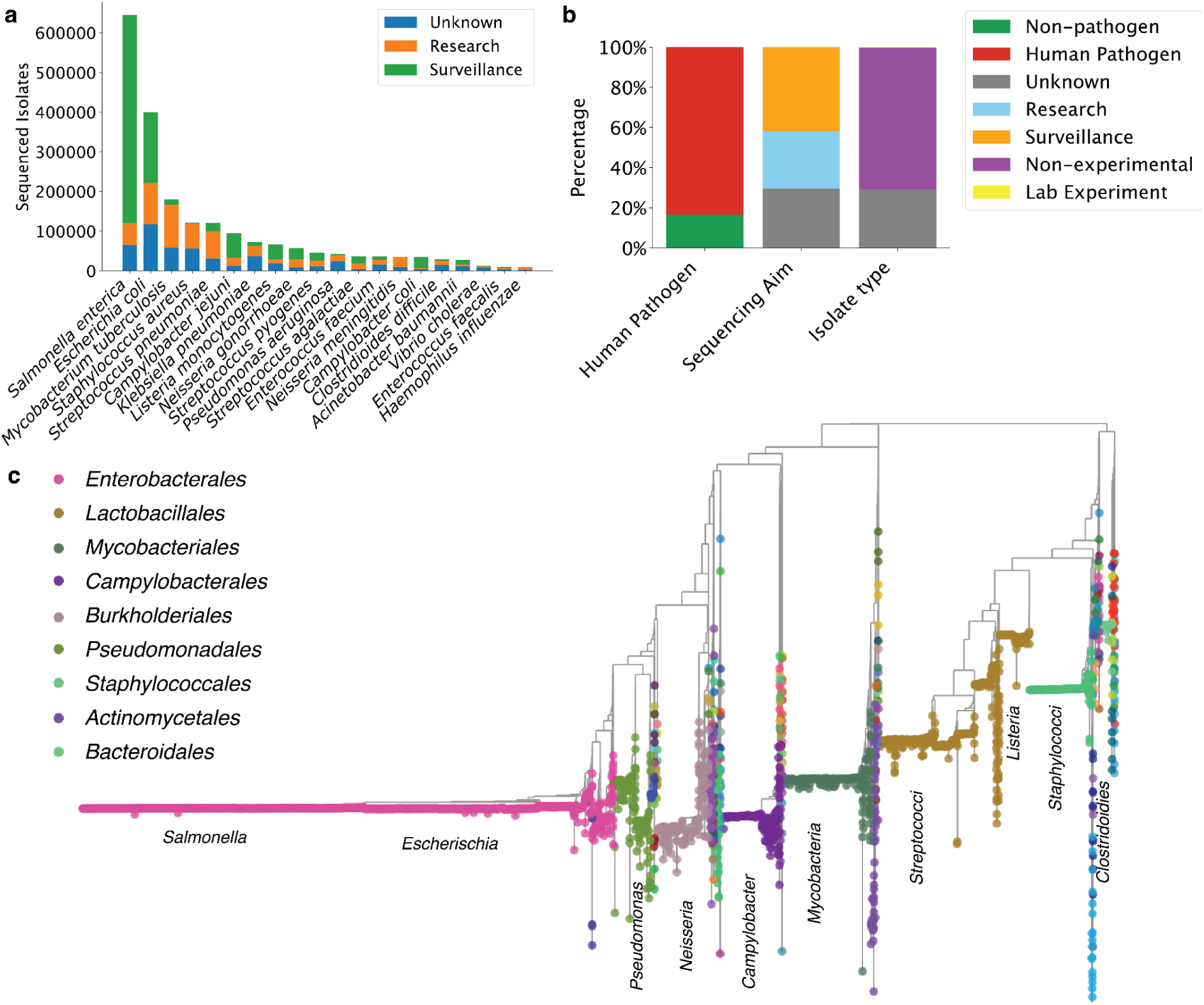
Composition and phylogenetic breadth of AllTheBacteria. a) Numbers of samples for the most common species, partitioned by whether sequenced for research or surveillance purposes (categorised at the study level), and a comparison with a previous dataset of 661,405 samples (Supp. Fig 2)^7^. b) Distribution of samples by pathogen status, study purpose and isolate type. (Lab Experimental: 0.27%) c) Hierarchical phylogeny of the high-quality portion of the dataset, inferred from 120 conserved marker genes^8^. Tree tips are coloured by order and major pathogen families or heavily sampled groups are labelled. The interactive tree is available at https://taxonium.org/atb

AllTheBacteria was co-created by a community of bioinformaticians, evolutionary biologists, public health researchers, and clinical microbiologists. Three principles guided the collaboration: unrestricted public release of derived data, reproducible software workflows and maximization of quality, utility and accessibility. The core pipeline performs genome assembly, taxonomic assignment and gene annotation, followed by community-prioritized analyses such as MLST, serotyping, antimicrobial-resistance profiling, protein-structure prediction for novel proteins, antiphage-defence detection and conversion of annotations into AI-ready formats (Supp. Fig 1). This design makes AllTheBacteria more than a collection of assemblies: it is a shared substrate on which multiple communities can build.

A major use case for AllTheBacteria is comprehensive search for genomes or sequence features of interest. For example, AllTheBacteria has already been used to search for an aminoglycoside resistance gene (npmA) across the entire dataset, expanding two hits to hundreds, allowing determination of host species, and mapping global spread^9^. Similarly, antibiotic resistance genes carried on plasmids and transposons have been tracked^10,11^; and emergence and prevalence of resistance within a species^12^. Beyond antimicrobial resistance, AllTheBacteria has supported mapping of Klebsiella autoinducers and their specificity^13^ and assessment of biosynthetic-gene-cluster conservation to aid natural-product discovery^14^. These examples illustrate how a harmonised public genome atlas converts isolated observations into population-scale biological context.

AllTheBacteria has also been used to address basic evolutionary questions. For example, showing that many apparent orphan genes do have homologues when one is able to scan across broader bacterial diversity, supporting horizontal transfer rather than de novo gene birth for a large fraction of cases^15^. It has also enabled generalization of a within-species model of deletion-generated gene fusions by screening similar events across Bacteria^16^. Evidence of selection in accessory genes is usually debated from a theoretical perspective, but our dataset was used to fit a model of accessory gene dynamics across bacteria, finding that the ratio of fast to slow moving accessory genes is the main difference between species pangenomes^17^. An analysis of per-gene %GC for genomes, using consistent criteria, identified this measure’s potential utility for genome quality control, as well as providing insight into an organism’s clonality or ecological isolation/specialization (Supp. Note 1, Supp. Figs. 3-6). These studies demonstrate that the value of the resource lies not only in its size, but in the ability to apply a single analytical framework across many taxa.

## Data volumes inspire new algorithms and AI-ready microbiology

The explosive growth of genomic data volumes posed considerable challenges for this project, and some central use-cases drove us to develop key algorithmic innovations. The first challenge was storage and distribution: gzip-compressed assemblies occupy terabytes, which would undermine the goal of broad accessibility. Because bacterial genomes are related by shared evolutionary history rather than random strings, we reasoned that compression would improve if genomes were batched by phylogenetic relatedness. This led to Miniphy^18^, which compresses 2.4 million genomes into 114 GB, making a resource of global scale practically distributable.

The second challenge was alignment. A core operation in genomics is comparing a query sequence, such as a gene or allele, with a genome collection while allowing evolutionary variation. At the scale of millions of genomes, existing approaches can require prohibitive memory or runtime. We therefore developed LexicMap^19^, an alignment method that searches millions of genomes in seconds to minutes, depending on hit number, with low memory use (∼10 GB RAM). This converts AllTheBacteria from a passive archive into an interactive search engine for microbial biology.

The third challenge was neighbour search and subset extraction. Many users need only the closest relatives of a focal genome or a defined subset of a species, not the entire collection. To support this, we created sketchlib.rust^20^, a scalable k-mer sketching tool, that combines efficient implementation, memory-mapping and streaming of data to reduce memory use. By integrating this into an AllTheBacteria command-line tool, this allows laptop-based scanning and subsetting of the entire dataset. As an example, we used it to contextualize a meningococcal outbreak in the United Kingdom in 2026^21,22^. Identifying 100 near-neighbour genomes took seven seconds, and phylogenetic analysis supported a single outbreak. Examination of the branch leading to the outbreak identified nine unique recombination events, two overlapping known antigens, PorB and pilus (Supp. Note 2, Supp. Figs 7,8). LexicMap searches then showed that these alleles had been present in other meningococcal genomes for at least two decades and were globally distributed, arguing against the outbreak being driven by a newly emerged, unusually transmissible genotype.

More broadly, AllTheBacteria provides benchmarks and data products that are already stimulating method development across bioinformatics and machine learning^23–26^. We also generated AlphaFold2 models^27^ for proteins without UniProt50 matches and deposited them as an AlphaFoldDB30 community dataset to facilitate structure-based search and reuse. We additionally provide genome annotations in an Apache Parquet dataset hosted on HuggingFace, containing gene and intergenic sequences, locations and genomic context for all samples (16.7 billion rows). Unlike full Bakta output, which is optimized for reproducibility, this representation is optimized for high-bandwidth access by AI training pipelines and has already supported transformer models of bacterial gene order^28,29^. Because oversampled lineages can distort model training, we further provide sketch-based deduplication so that users can select representatives at chosen identity thresholds. These design choices make AllTheBacteria a scalable foundation for data-driven microbiology.

## AMR reservoirs beyond priority pathogens

Bacterial use of antimicrobials long predates human industrialisation of antibiotic use^30^, so the distribution of antimicrobial resistance (AMR) genes across species and niches provides insight into both clinical risk and environmental reservoirs. AMR surveillance tends to focus on key opportunistic pathogens which bear a high AMR load: *Enterococcus faecium*, *Staphylococcus aureus*, *Klebsiella pneumoniae*, *Acinetobacter baumannii*, *Pseudomonas aeruginosa*, *Enterobacter* spp., and *Escherichia coli* (referred to by the acronym ESKAPEE). Metagenomic sequencing can identify environments enriched for resistance genes, but it generally cannot assign those genes to isolate-resolved host species. We used AllTheBacteria to scan 1,087 non-priority species for known AMR determinants. Outside the ESKAPEE pathogens, AMR burden varied markedly by genus (Fig. 2a). Most rare genera (17 of 27; 53%; defined in Methods) carried only one to two known AMR genes per genome, whereas ESKAPEE genera showed median burdens from 5 in *Escherichia* to 15 in *Klebsiella* and *Enterococcus* (Fig. 2b). However, selected non-priority genera, including *Mannheimia*, *Citrobacter*, *Bacillus* and *Vibrio*, carried higher median burdens (7, 5, 4 and 4 genes, respectively), and *Proteus*, *Vibrio*, *Citrobacter*, *Serratia* and *Aeromonas* showed broad ranges with maxima of 26-32 genes. Linking these genera to biome annotations from SPIRE^31^ revealed that human-associated sampling dominates the public record, but also highlighted water-associated genera with contrasting AMR profiles, such as *Vibrio* and *Legionella*. Thus, AllTheBacteria enables isolate-resolved AMR reservoir discovery beyond the pathogens that dominate surveillance.

**Figure 2.**
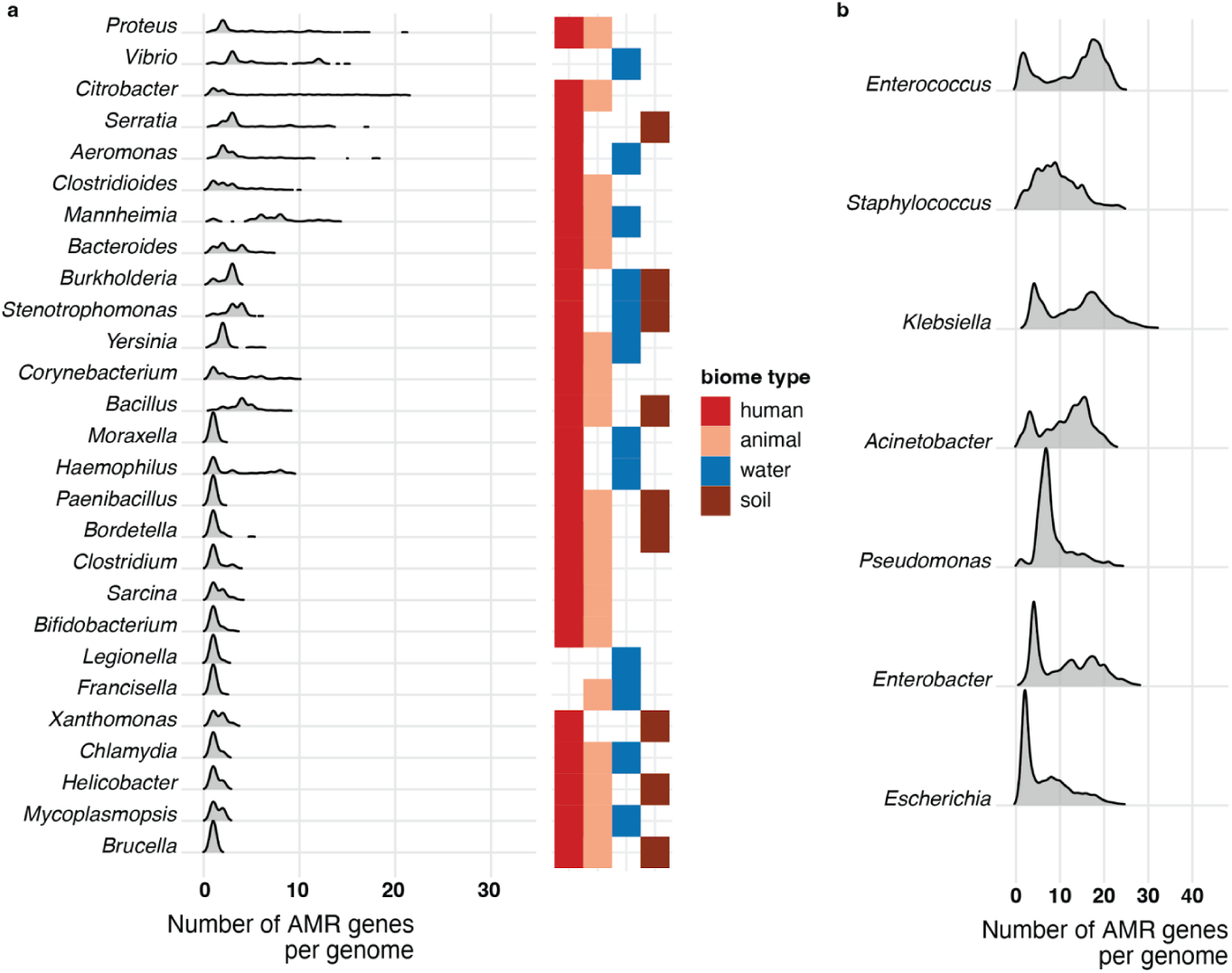
AMR genes across genera, and genera across biomes. a) (left) Distribution of number of AMR genes across multiple genera excluding ESKAPEE genera and the top 10 most common in AllTheBacteria; (right) presence of the genus in human, animal, water and soil biomes as detected in SPIRE. b) For comparison the distribution of acquirable AMR genes in the ESKAPEE genera in AllTheBacteria, all major pathogens.

## Antiphage defense element prevalence and interactions across Enterobacterales

We next used AllTheBacteria to examine antiphage-defence systems across 27 Enterobacterales species with at least 400 genomes each (n = 1,166,956 genomes). We identified 226 unique defence systems. Most were rare (Supp. Fig 9), but 22 were detected at least once in every species analysed (Fig. 3). Restriction-modification was the only system present at high within-species frequency across all species. The toxin-antitoxin systems Mok-Hok-Sok and MazEF were also abundant^32,33^. By contrast, systems such as BstA, Gao_let, PrrC, Rst_3HP, Gao_Tmn and Shedu were broadly distributed across species but always at low within-species frequency, suggesting niche-specific benefits, costs of maintenance or fluctuating advantage under high gene turnover^34^. CRISPR-Cas systems were present in 26 of 27 species, with *Salmonella arizonae* as the exception^35^. Interaction analysis with SimPhyNI^36^ identified 39 significant interactions between 15 systems across the full phylogeny (Benjamini-Yekutieli-corrected P < 0.05), 90% of which were avoidances. Within *E. coli*, however, 101 significant interactions were split almost equally between avoidance and co-occurrence, including positive interactions between systems with experimentally demonstrated synergy such as Zorya II and Druantia III^37^. These results show that AllTheBacteria enables exploration of defence-system ecology at both pan-species and within-species scales.

**Figure 3.**
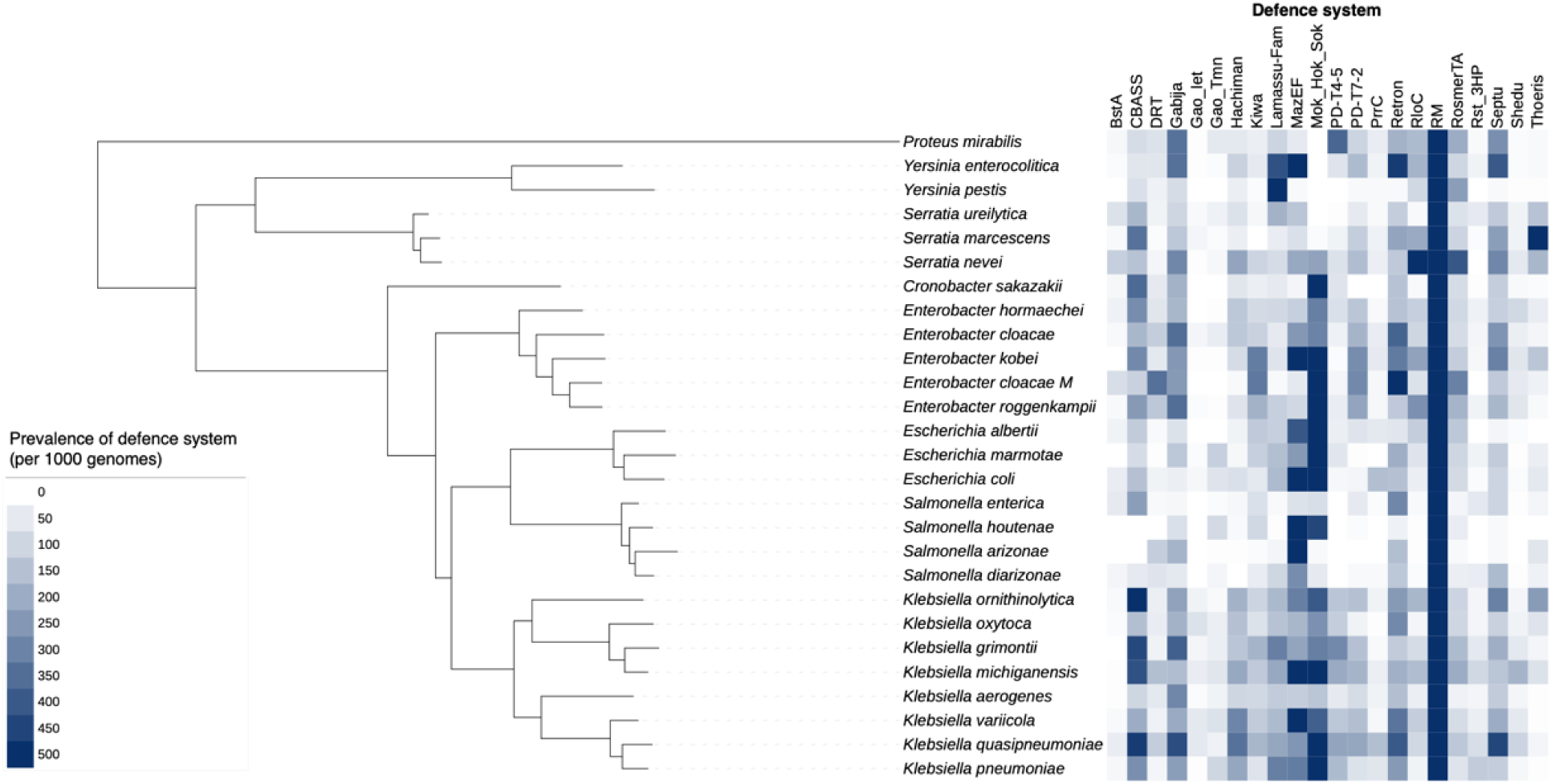
Within-species frequency of defence systems found across Enterobacterales. (Left) Phylogeny of species within this analysis. (Right) Within-species frequency (shown by colour) of defence systems (columns) in each species.

## AI mining of AllTheBacteria reveals encrypted peptide antibiotics

To test whether AllTheBacteria can directly generate experimentally tractable biological discoveries, we mined bacterial proteomes for encrypted peptides: short protein fragments whose antimicrobial activity is not obvious from the parent protein but can be predicted from sequence and physicochemical features^38^. We screened 3,919,096 short peptide fragments (<50 amino acid residues) with APEX 1.1^39^, a deep learning model that predicts peptide antimicrobial activity across 11 bacterial pathogens. This yielded 1,867 candidates with predicted minimum inhibitory concentration (MIC) values below 80 μmol L^−1^ against at least one pathogen, creating a focused set of putative antimicrobial molecules from a proteome-scale search space.

We first compared the predicted AllTheBacteria (here abbreviated ATB) peptides with antimicrobial peptides curated in widely used databases (see Supp. Note 3, Supp. Figs 10-13). ATB peptides were enriched in basic residues, particularly lysine and arginine, and many also contained hydrophobic or aromatic residues, including leucine, isoleucine, and tryptophan. These features are consistent with electrostatic engagement of bacterial envelopes and potential membrane interaction. From this prioritized pool, we selected 24 peptides (Supp. Table 1) for chemical synthesis and experimental validation. The selected panel included highly cationic low-complexity sequences, mixed hydrophobic–cationic architectures and tryptophan-rich motifs, allowing us to test whether distinct classes of ATB-derived encrypted peptides translate into antibacterial activity. The 24 synthesized peptides were profiled against 20 clinically relevant Gram-negative and Gram-positive bacterial strains, including antibiotic-resistant ones (Fig. 4). Antibacterial activity was most pronounced against Gram-negative species, including *Acinetobacter baumannii*, *Escherichia coli*, *Enterobacter cloacae*, *Klebsiella pneumoniae*, and *Pseudomonas aeruginosa*.

**Figure 4.**
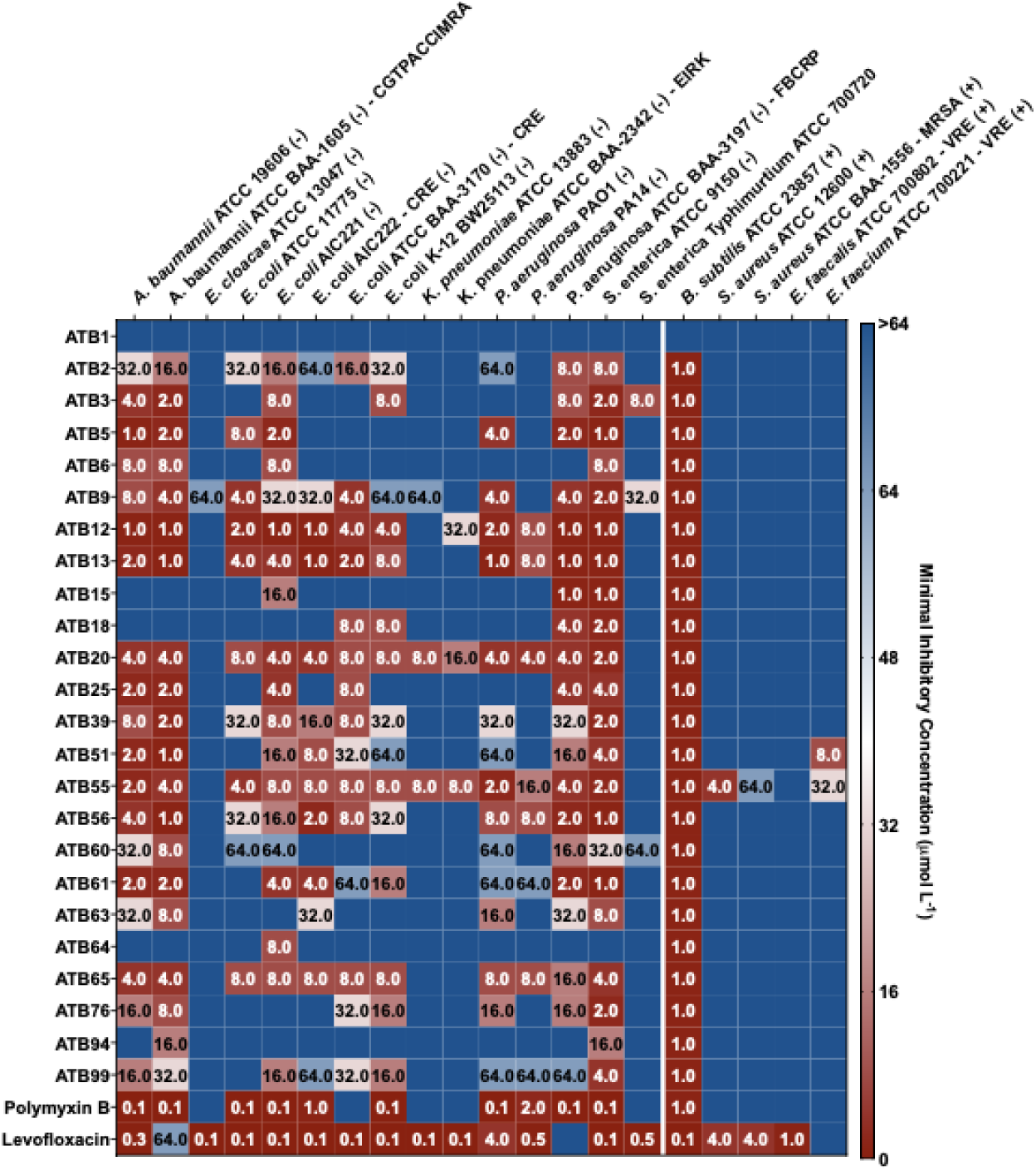
Antimicrobial activity of APEX-predicted AllTheBacteria peptides. Heat map displaying the antimicrobial activities (μmol L^−1^) of ATB peptides (rows) against 20 clinically relevant pathogens (columns), including antibiotic-resistant strains. Briefly, 10^5^ bacterial cells were incubated with serially diluted peptides (0–64 μmol L^−1^) at 37 °C. Bacterial growth was assessed by measuring the optical density at 600 nm in a microplate reader one day post-treatment. The MIC values presented in the heat map represent the mode of the replicates for each condition and the antibiotics polymyxin B and levofloxacin were used as positive controls.

Several peptides inhibited one or more strains at low micromolar concentrations. ATB5, ATB3, and ATB12 were particularly active against *E. coli* isolates, including a colistin-resistant strain, with MIC values of 2–4 μmol L^−1^. ATB5 also inhibited *A. baumannii* ATCC 19606 at 1 μmol L^−1^, placing it among the most potent peptides in the panel. ATB9, a tryptophan-rich, highly cationic polymeric motif, displayed a similarly broad profile, suppressing *E. coli* at 4 μmol L^−1^and *A. baumannii* at 8–64 μmol L^−1^ depending on the strain background.

Activity against Gram-positive pathogens was more limited but detectable for selected peptides. ATB51 inhibited methicillin-resistant *Staphylococcus aureus* (MRSA) at 8 μmol L^−1^, while ATB55 inhibited both MRSA and *S. aureus* ATCC 12600 at 32 μmol L^−1^. Several ATB peptides also retained activity against resistant Gram-negative organisms, including colistin-resistant *E. coli*, carbapenem-resistant *K. pneumoniae* and multidrug-resistant *P. aeruginosa*. Across the panel, activity was not explained by positive charge alone: highly repetitive cationic peptides such as ATB1 and ATB15 showed limited measurable activity, whereas peptides combining cationic residues with hydrophobic or aromatic content showed stronger antibacterial profiles. These data establish bacterial proteomes as a source of experimentally validated antimicrobial scaffolds and demonstrate that AllTheBacteria can be mined as a functional-molecule discovery engine.

## ATB peptides show membrane-responsive conformational transitions and peptide specific membrane perturbation

To assess whether ATB peptides undergo environment-dependent structural changes, we analysed their secondary-structure propensities by circular dichroism (CD) spectroscopy and quantified the spectra using BeStSel algorithm^40^. Peptides were measured in water, 50% methanol/water, 60% trifluoroethanol/water, and 10 mmol L^−1^ Sodium dodecyl sulfate, spanning aqueous to membrane-mimetic environments (Supp. Note 4, Supp. Fig 14). Across the panel, ATB peptides were largely disordered in aqueous solution but showed increased helical character in structure-promoting or membrane-mimetic conditions, particularly TFE and SDS. This behaviour is consistent with folding-upon-binding transitions observed for many membrane-active peptides and supports a link between ATB sequence composition, inducible structure and antibacterial activity.

We next tested whether ATB peptides act primarily by disrupting the bacterial membrane, we assessed their ability to depolarize the cytoplasmic membrane and permeabilize the outer membrane of *A. baumannii* ATCC 19606 at their MICs using DiSC_3_-5 and NPN uptake assays, respectively. In both assays, bacteria in buffer with dye served as the baseline condition, Triton X-100 was used as a non-specific membrane-disrupting positive control, and polymyxin B and levofloxacin were included as reference antibiotics. Fluorescence data were analysed in terms of both maximal signal over time and time-resolved kinetic profiles.

In the DiSC_3_-5 depolarization assay, a subset of peptides, including ATB39, ATB63, ATB60 and ATB65, generated larger depolarization signals than the remaining ATB peptides and antibiotic comparators, but these responses remained below the magnitude observed with the positive control, Triton X-100. These data suggest that rapid collapse of cytoplasmic membrane potential is not the dominant early response for most ATB peptides under these conditions.

The NPN uptake experiments then provided a distinct view of the outer membrane. Several ATB peptides produced sustained increases in NPN fluorescence above the bacterial baseline, whereas polymyxin B and levofloxacin remained at or near baseline in this assay. ATB76, ATB63, ATB60, ATB65, ATB61 and ATB99 generated the largest NPN responses, followed by ATB51, ATB39, ATB20, ATB55, ATB56 and ATB6. ATB2 and ATB5 produced only minor NPN increases. Thus, outer-membrane permeabilization is a prominent feature for a subset of ATB peptides, whereas cytoplasmic membrane depolarization is generally more modest.

Cytotoxicity profiling in human embryonic kidney (HEK293T) cells (see Supp. Note 5) revealed a spectrum of safety profiles within the ATB panel. Several peptides combined low-micromolar antibacterial activity with favourable selectivity indices, including ATB20, ATB51, ATB9, ATB61, ATB12 and ATB2. In contrast, ATB25, ATB55, ATB60, ATB65 and ATB76 showed narrower therapeutic windows, indicating that some membrane-active ATB peptides may require chemical optimization to reduce mammalian-cell toxicity. Together, the structural, membrane-perturbation and cytotoxicity data indicate that ATB peptides occupy a range of membrane-active behaviours, with the most promising leads balancing antibacterial potency with acceptable selectivity.

## Anti-infective activity of an ATB peptide in mouse models

We next asked whether an AllTheBacteria-derived peptide could reduce bacterial burden *in vivo*. We selected ATB20 because it combined antibacterial potency, favourable selectivity in human kidney cells and membrane-targeting behaviour. We tested it in a murine *A. baumannii* ATCC 19606 skin abscess model^41–43^ (**Fig. 5a**), a clinically relevant infection setting for a pathogen associated with skin, soft-tissue and deep-tissue infections, as well as multidrug-resistant hospital outbreaks.

**Figure 5.**
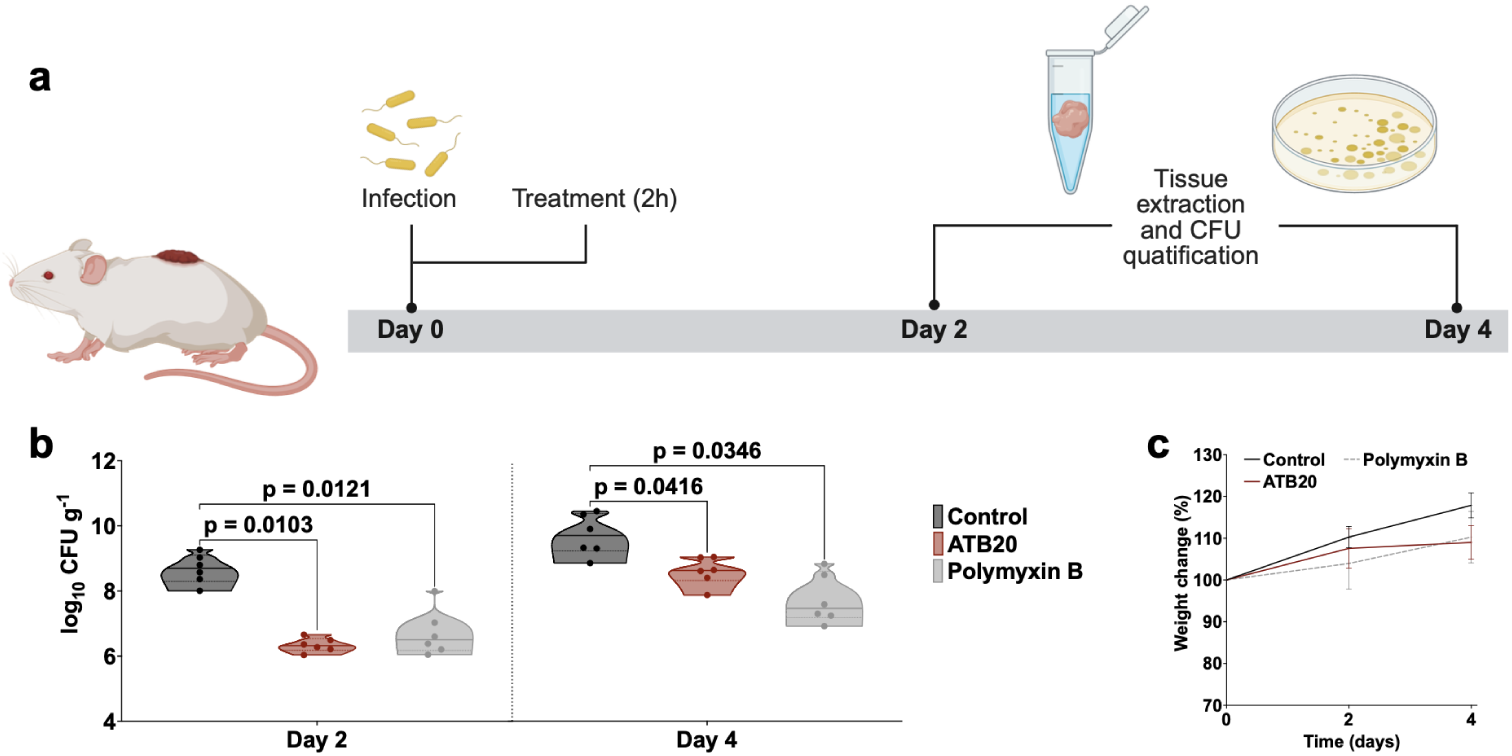
Anti-infective activity of a lead ATB peptide in a murine infection model. **(a)** Schematic representation of the skin abscess mouse model used to assess the anti-infective activity of ATB20 (n = 6) against *A. baumannii* ATCC 19606. **(b)** Level of infection (in log of colony forming units (CFU) per gram) in control group, and treatment with either ATB20 or Polymyxin B. ATB20 administered at 10xMIC in a single dose post-infection, inhibited the proliferation of the infection for up to 4 days after treatment compared to the untreated control group. ATB20 reduced the infection by up to 2.5 orders of magnitude (day 2), demonstrating activity comparable to the control antibiotic, polymyxin B. **(c)** Mouse weight was monitored throughout the duration of the skin abscess model (4 days total) to assess potential toxic effects of both the bacterial load and ATB20. The limit of detection (LOD) for the CFU quantification is log_10_ CFU = 2. Bacterial burdens are shown as violin plots with individual data points overlaid; each point represents one animal. The horizontal line within each violin indicates the mean, and violin width reflects the distribution of values within each group. Statistical comparisons were performed separately for each time point using ordinary one-way ANOVA followed by Dunnett’s two-sided multiple comparisons test versus the control group. Exact p-values are shown above the indicated comparisons. (Panel **a** created in BioRender.)

A single topical dose of ATB20 at 10×MIC reduced bacterial burden by ∼2-log two days after infection relative to untreated controls, with efficacy comparable to polymyxin B (**Fig. 5b**). By day 4, ATB20 maintained ∼1.5-log reduction and again matched polymyxin B. Mouse body weight remained stable throughout the experiment, indicating no overt treatment-associated toxicity under these conditions (**Fig. 5c**). These results move AllTheBacteria-derived peptides beyond in vitro activity and establish ATB20 as an in vivo-active antimicrobial lead.

## Discussion

High-quality open datasets accelerate biology by reducing duplicated effort, allowing individual researchers and smaller laboratories to analyse data they could not generate alone, centralizing quality control and creating foundations on which new methods can be built. Human genetics has repeatedly demonstrated this effect through HapMap, the 1000 Genomes Project, ExAC and gnomAD, which turned aggregated variation data into community standards for discovery and benchmarking. Microbiology has lacked an equivalent because public bacterial sequencing has grown faster than reusable infrastructure. AllTheBacteria addresses this gap by combining standardized assembly, quality control, taxonomy, annotation, searchable indexes and community governance. Its adoption as a benchmark, search resource and source of contextual genomes reflects the requirement for not only more data, but better organized data.

The antimicrobial discoveries reported here illustrate why scale and standardization matter. By coupling AllTheBacteria with APEX, we moved from millions of proteome-derived peptide fragments to a small experimentally testable set and identified multiple peptides with low-micromolar activity, mechanistic evidence of membrane interaction and in vivo efficacy for ATB20. The ATB-derived peptides overlap with known antimicrobial design principles but also occupy distinct physicochemical space, with high cationicity, compact hydrophobic patterning and membrane-responsive folding. This suggests that bacterial proteomes encode antimicrobial-like fragments that are not fully represented in existing peptide databases. The same infrastructure can be linked to external phenotype resources such as CABBAGE^44^ and EMBL-EBI’s AMR portal^45^, extending its use toward AMR prediction, diagnostic development and therapeutic discovery.

AllTheBacteria is designed to remain current. After the initial 2.4-million-assembly release covering data up to August 2024, we released an additional 320,000 samples (2.8 million total, up to May 2025) and are processing a further 350,000 samples (3.1 million total, up to March 2026). Continued maintenance will require sustained community effort and funding, but the software workflows are standardised and reproducible, and the processing model is deliberately distributed: core assembly, quality control and sharing remain centralized, while specialized analyses can be contributed by communities focused on particular genera, mobile elements, molecular functions or applications.

The recent revolution in protein-structure prediction shows what becomes possible when modern AI is paired with decades of harmonized biological data, trusted benchmarks and active communities^27,46^. AllTheBacteria provides an analogous substrate for microbial genomics: a standardized, openly reusable and continually extensible representation of public bacterial and archaeal diversity. By showing that this resource can support outbreak analysis, resistance surveillance, defence-system ecology, AI method development and the discovery of in vivo-active peptide antibiotics, we establish AllTheBacteria as infrastructure for a new generation of data-driven microbiology.

## Methods

### Prior dataset and process on which AllTheBacteria was built

Blackwell et al. (2021) set out to uniformly assemble, QC and analyse all bacterial isolate whole genome sequence (WGS) raw data available in the European Nucleotide Archive (ENA) as of November 2018^7^. They released 639,981 high-quality assemblies, along with quality control information and fundamental genome-derived statistics – the most important of which was to check the taxonomic abundance (using the NCBI taxonomy) within each putatively single isolate dataset to confirm the species label in the submitted ENA metadata, which is not necessarily sequence-derived. In the process they estimated that 8.1% of the species metadata tags in the ENA were incorrect. We refer to this dataset as the 661k dataset.

### Dataset

We downloaded all paired Illumina bacterial isolate whole genome sequence raw sequence metadata from the ENA, using the query https://www.ebi.ac.uk/ena/portal/api/search?result=read_run&fields=ALL&query=tax_tree(2)&format=tsv

Samples were processed if they were not in the 661k dataset, and had metadata instrument_platform=ILLUMINA, library_strategy==WGS, library_source= GENOMIC, and library_layout=PAIRED. The samples were processed in two stages: the first (release 0.2) was from metadata downloaded on June 16th 2023, and then a second round of processing from metadata obtained on August 1st 2024 (incremental release 2024-08).

To understand the provenance of the isolates included in the dataset, we classified isolates according to their ENA study accessions in two ways. First, we examined the project name, description, submitting center, and associated publications of all studies with 500 or more isolates. Projects sequenced by public health laboratories with no described research purpose or associated publications were classified as genomic surveillance isolates. Projects sequenced by research institutions with explicit research aims were classified as research. This includes surveillance ‘pilot’ projects which were performed to investigate or demonstrate the potential of genomic surveillance. Project descriptions and associated publications were also consulted to determine if the sequenced isolates were the result of experimental work. In both the case of surveillance and experimental classification, studies without sufficient metadata were not classified.

All isolates included in the study were also classified as human pathogens (or not) based on a previously published comprehensive list of pathogens^47^.

### Software and pipelines

Reproducibility of all bioinformatics processing was ensured in two ways. First, all code used for processing is maintained in github. Processes that are run on all samples are here: https://github.com/AllTheBacteria/AllTheBacteria/tree/main/reproducibility/All-samples and species-specific pipelines here: https://github.com/AllTheBacteria/AllTheBacteria/tree/main/reproducibility. There is a process for contributing to the project, documented here https://allthebacteria.org/docs/contributing/, which involves another person confirming that there is code for reproducibility, and documentation.

### Genome assembly

The genome assembly pipeline for the 661k dataset used by Blackwell was based around v1.0.4 of Shovill (https://github.com/tseemann/shovill) which is a wrapper around Spades^48^. For release 0.2, we refactored and updated the pipeline (https://github.com/leoisl/bacterial_assembly_pipeline), and used an updated version of Shovill (v1.1.0), however the difference between v1.0.4 and v1.1.0 was minimal and does not impact assembler output, and therefore there was no need to reassemble the existing 661k dataset.

All samples in release 2024-08 were processed using a simple Python script (see https://github.com/AllTheBacteria/AllTheBacteria/tree/main/reproducibility/All-samples/assembly), again using Shovill v1.1.0. The script processes one sample, first downloading the reads, then running Sylph, Shovill, and finally removing contigs matching the human genome (as described later).

### Taxonomic abundance estimation

Most taxonomic abundance estimation tools are designed for metagenome data which consists of an unknown mix of different taxa. However, in single isolate data, which makes up the entirety of this collection, only a single species is expected to be present, unless the sample is contaminated. Therefore performing taxonomic analysis on isolate data is considerably simpler than on full metagenomic data. We wanted primarily to establish the major species, its relative abundance, and the nature of contaminants. We used sylph^49^ version 0.5.1 with the pre-built GTDB r214 database (https://storage.googleapis.com/sylph-stuff/v0.3-c200-gtdb-r214.syldb) with default options, which required just 10Gb of RAM and took (∼1 minute per sample). Since the 661k dataset had previously been analysed with Kraken/Bracken, we re-downloaded the reads and reprocessed them with sylph.

A species call was made from the ‘Genome_file’ column of the sylph output, using a lookup table generated with TaxonKit^50^ using GTDB taxonomy data (https://github.com/shenwei356/gtdb-taxdump v0.4.0). The reads from 3,252 samples resulted in no output from sylph, presumably because there were no matches to the reference database.

### Human decontamination

After assembly, the contigs output by Shovill were matched to the human genome plus HLA sequences using nucmer from version 4.0.0rc1 of the MUMmer package^51^. We used the T2T CHM13 version 2 assembly (GCA_009914755.4) of the human genome^52,53^. For HLA sequences, we used the file hla_gen.fasta from version 3.55.0 of the IPD-IMGT/HLA database^54–56^. Any contig that had a single match of at least 99% identity and 90% of its length was removed.

### Assembly statistics

The program assembly-stats (https://github.com/sanger-pathogens/assembly-stats; git commit 7bdb58b) was run on each assembly to gather basic statistics. Assemblies with a total length of less than 100kbp or greater than 15Mbp were excluded. We found that 21 of the assemblies in the original 661k data set were longer than 15Mbp, and so were removed from our releases, meaning that 661,384 of the samples in AllTheBacteria originate from the 661k dataset.

### CheckM

CheckM2^57^ version 1.0.1 was run on each assembly, using the default downloaded database uniref100.KO.1.dmnd. We ran *checkm2 predict* with options --allmodels --database_path --lowmem. We found 275 samples did not run to completion, stopping with the error message “No DIAMOND annotation was generated”. This suggested that those assemblies were of low quality, resulting in very few predicted proteins.

### MiniPhy

All assembly FASTA files were compressed using MiniPhy^18^ commit 7abe08c, which uses intelligent batching of genomes to improve compression. The process has two steps:

1. Divide the genomes into approximately equal-sized batches, typically done by species. In our case, the highest-abundance species for each sample was previously determined using sylph (see above), and a CSV file was created mapping the filename to species. Batches were auto-created using the create_batches.py script from the MiniPhy repository.
2. MiniPhy was then run on each batch; internally it created an approximate phylogenetic tree and reordered the genomes for better compression. The output is then compressed with the standard xz tool, to produce one archive file per batch.

This results in compressed archive files of total size 114 GB holding 2,440,377 genomes; for comparison the full data would take 4 Terabytes of space if each assembly was individually gzipped.

### sketchlib.rust

pp-sketchlib is a tool for rapid sketching and distance estimation of genomes which scales well to datasets of millions of genomes^20,58^ (orders of magnitude faster than mash/sourmash^20^). We reimplemented the tool in Rust, using a new dataset format for storing and accessing^20^ sketches which scales linearly in access time for the number of sketches being accessed. The high-quality assemblies were sketched at k=17 using sketchlib.rust v0.1.0. This database allows sequence similarity search through computing a Jaccard index, either against all the contents, sparse queries returning k-nearest neighbours, below a given distance threshold, or against a chosen subset of queries. We ran sketchlib sketch -f 2kk_list.txt -k 17 -s 1024 -o 2kk_sketch --threads 32 to use a sketch size of 1024. The resulting .skd database of sketches is 4.1 GB, and .skm of metadata is 123 MB.

### Phylogeny construction

An alignment was constructed using 120 bacterial marker genes from GTDB (https://www.nature.com/articles/nbt.4229); each sample was annotated with pyrodigal (https://pyrodigal.readthedocs.io/en/stable/), and then pyhmmer (https://pyhmmer.readthedocs.io/en/stable) was used to align the predicted genes with the profile HMMs for each marker gene. Alignments were trimmed to the length of their respective marker gene profile HMM and were concatenated to create a global alignment for all samples. This broadly follows the method described here: https://pubmed.ncbi.nlm.nih.gov/16513982/.

The tree was built by a divide-and-conquer approach. Representatives were picked using gemsparcl (https://www.biorxiv.org/content/10.64898/2025.12.30.695181v1.abstract), resulting in 21,593 top-level clusters. These were used to build a backbone tree. For each cluster with at least three samples, a tree was built within the cluster, including its nearest neighbour in the backbone tree as an outgroup. These subtrees were then grafted to the backbone tree using the same root between the outgroup as in the subtree and backbone tree. Small trees were built with IQtree v3.0.1 (https://academic.oup.com/mbe/article/43/5/msag117/8669857) in fast mode. The seven largest trees were built using VeryFastTree v4.0.5 (https://doi.org/10.1093/bioinformatics/btaa582). All trees were built using the LG amino acid substitution matrix (https://pubmed.ncbi.nlm.nih.gov/18367465/) with invariant sites and gamma distributed rate heterogeneity with four rate compartments. All scripts used for tree construction are available on GitHub (https://github.com/bacpop/ATB_tree)

Each cluster was assigned to a species using GTDB-tk v2.4.1 (https://academic.oup.com/bioinformatics/article/36/6/1925/5626182) using GTDB release 226 (https://academic.oup.com/nar/article/54/D1/D743/8296754). The full taxonomic designation was filled out using TaxonKit (https://bioinf.shenwei.me/taxonkit/tutorial/). These were then copied across all members of each cluster.

### Defence System detection

We ran DefenseFinder v1.3.0 (https://github.com/mdmparis/defense-finder; models version v.1.3.0 CasFinder v3.1.0) on all available Enterobacterales genomes for species within the ATB database with 400 or more genomes, this equated to a total of 1,166,956 genomes. Note: this is all GTDB species with >400 samples. This comprises 27 species: *Salmonella enterica, Escherichia coli, Klebsiella pneumoniae, Enterobacter hormaechei A, Salmonella houtenae, Salmonella diarizonae, Klebsiella quasipneumoniae, Yersinia enterocolitica, Proteus mirabilis, Klebsiella variicola, Yersinia pestis, Salmonella arizonae, Klebsiella michiganensis, Cronobacter sakazakii, Klebsiella aerogenes, Serratia marcescens K, Klebsiella oxytoca, Enterobacter kobei, Serratia nevei, Enterobacter roggenkampii, Escherichia albertii, Enterobacter cloacae, Klebsiella ornithinolytica, Klebsiella grimontii, Escherichia marmotae, Enterobacter cloacae M, Serratia ureilytica*.

To validate the effectiveness of DefenseFinder on the AllTheBacteria Illumina assemblies, we compared results within *S. enterica* for RefSeq complete assemblies and AllTheBacteria assemblies. The average number of systems per genome was only slightly lower in AllTheBacteria compared to RefSeq (7.05 vs. 7.91) and system prevalences were highly correlated (Pearson’s R=0.90; Supp. Fig 15) suggesting that using DefenseFinder within AllTheBacteria assemblies is consistent with using it on only complete genomes.

### Defence system association and avoidance testing

We randomly subsampled both the full phylogeny and the *E. coli* subtree to 100,000 tips. We then ran SimPhyNI (v. 1.0.2) with default parameters, considering defence system subtypes between 5% and 95% prevalence. The script to generate the subsampled trees is available at https://github.com/wtmatlock/atb-defensefinder-simphyni, and the output of SymPhyNI is here https://github.com/AllTheBacteria/atb-paper/tree/main/analysis/defence_interactions. Across the full diversity phylogeny, the prevalence of negative associations likely reflects SimPhyNI’s reliance on convergent signals (repeated, independent transitions). This creates an asymmetric burden of proof where recurrent mutual exclusion readily produces independent events, whereas lineage-restricted positive associations are often attributed to shared ancestry and therefore discounted as evidence of interaction. In *E. coli*, however, high rates of horizontal gene transfer likely generate frequent co-acquisition events, providing the repeated independent transitions necessary for SimPhyNI to statistically distinguish convergent association from vertical inheritance of system pairs.

### Antimicrobial resistance gene and mutation detection

Antimicrobial resistance determinants were identified using AMRFinderPlus^59^ v3.12.8 on all assembly FASTA files with database version v2024-01-31.1. For appropriate species (*Acinetobacter baumannii, Burkholderia cepacia, Burkholderia pseudomallei, Campylobacter jejuni, Campylobacter coli, Citrobacter freundii, Clostridioides difficile, Enterobacter cloacae, Enterobacter asburiae, Enterococcus faecalis, Enterococcus faecium, Enterococcus hirae, Escherichia, Shigella, Klebsiella aerogenes, Klebsiella oxytoca, Klebsiella pneumoniae, Klebsiella pneumoniae species complex, Neisseria gonorrhoeae, Neisseria meningitidis, Pseudomonas aeruginosa, Salmonella, Serratia marcescens, Staphylococcus aureus, Staphylococcus pseudintermedius, Streptococcus agalactiae, Streptococcus mitis, Streptococcus pneumoniae, Streptococcus pyogenes, Vibrio cholerae, Vibrio vulnificus, Vibrio parahaemolyticus*) we used the GTDB species assigned by sylph for the AMRFinderPlus *--organism* parameter, in accordance with the guidelines at https://github.com/ncbi/amr/wiki/Running-AMRFinderPlus#--organism-option (git commit 5f27bbe), thus incorporating known AMR-informative point mutations. This option was omitted for all other species.

We extracted AMRFinderPlus results for all genera with at least 1000 genomes. ‘Rare’ genera were defined as those that i) weren’t in the top 10 of genera by genome counts (i.e. excluding *Salmonella, Escherichia, Streptococcus, Mycobacterium, Staphylococcus, Campylobacter, Neisseria, Klebsiella, Listeria, Enterococcus*), and ii) didn’t belong to the ESKAPEE pathogens (this excludes *Pseudomonas*, *Acinetobacter* and *Enterobacter*). We summed the total number of AMR determinants per genome, excluding those labelled as “EFFLUX”. For each rare genus, we extracted the top 3 biomes they were found in from the SPIRE^31^ study. Each biome was classified as either human, animal, water or soil.

### Multi-locus sequence typing

Multi-locus sequence typing (MLST) was conducted using MLST v2.32.2 (https://github.com/tseemann/mlst) and MLSTdb v1.3.0 (https://github.com/MDU-PHL/mlstdb) against all schemes available within the PubMLST^60^ and PasteurMLST databases from the date of 2026-02-03 for a total of 167 schemes. MLST schemes were automatically assigned based on the best matching allele profiles with the ecoli, abaumannii, vcholerae_2 schemes excluded from autodetection. In a number of instances, genomes produce multiple equally strong “PERFECT” matches, resulting in ambiguous assignments for which both results were recorded. From this list, we identified a subset of 47 profiles belonging to five schemes that were considered to be cross-species contaminants. For example, ecoli-achtman_4(ST102) was not found in a single *Escherichia coli* genome and was exclusively found in *Salmonella enterica* genomes.

### Virulence factor profiling

Virulence gene screening was performed using Abricate v1.0.1 (https://github.com/tseemann/abricate) against the Virulence Factor Database (VFDB, 2024 release)^61,62^. Gene matches were reported using minimum thresholds of 80% nucleotide identity and 80% coverage. Results were generated for all genomes, including those without detectable virulence factor matches.

### Biosynthetic gene cluster detection

Biosynthetic gene clusters (BGCs) were detected in each genome using GECCO^63^ v0.9.8 (executed via Nextflow^64^ v24.04.2). Briefly, each assembled genome was supplied as input (in FASTA format) to the “gecco run” command, which was used to detect BGCs with the “--mask” and “--merge-gbk” options enabled (remaining parameters were set to their defaults). The resulting BGC-related output files (i.e., BGC GenBank files and BGC cluster TSVs) were concatenated and stored.

### Species-specific genotyping

Using sylph-identified species designations the following bioinformatics tools were used to provide additional species-specific genotyping.

Briefly *Acinetobacter baumannii* capsule types O and K^65^ were assigned with Kaptive3^66,67^. *Escherichia coli* capsule types were assigned using ECtyper^68^ v2.0.0, using the –verify and –pathotype flags to enable serotype, pathotype and species classification. A subset of genomes identified by Sylph as *Escherichia coli* and belonging to MLST sequence types known to contain *Shigella* spp. within the ecoli-achtman_4 MLST scheme were further analysed using ShigEiFinder^69^ using default parameters to assign *Shigella* serotypes and *EIEC E.coli* pathotypes. *Haemophillus spp* polysaccharide capsule types were assigned using *hicap*^70^ *v1.0.4* using default parameters*. Klebsiella spp* were additionally typed with kleborate^71^ v2.3.3 and kaptive3^66^ providing speciation, virulence gene detection, MLST, capsule and antimicrobial resistance profiles, run with default parameters. *Legionella pneumophila* sequence based typing scheme was assigned with legsta v0.5.1 (https://github.com/tseemann/legsta) using default parameters. Listeria monocytogenes serotypes were assigned using LisSero^72^ v0.4.9 with minid and mincov parameters set at 95. *Mycobacterium tuberculosis* were typed with TBProfiler^73^ v6.2.1 providing spoligotype, lineage and antimicrobial resistance profiles. *Neisseria gonorrhoeae* were typed with pyngoST^74^ v1.1.2 with the CC database 240513_NGSTAR_CC_updated to provide multi-antigen sequence typing (NG-MAST) and sequencing typing for antimicrobial resistance (NG-STAR) and penA allele results. Neisseria meningitidis were typed using meningotype^75^ v0.8.5 (https://github.com/MDU-PHL/meningotype) which provided serotype, MLST, Finetyping^76^, Bexsero antigen typing^77^, and MendeVAR^78^ (meningococcal Deduced Vaccine Antigen Reactivity Index) results. *Streptococcus algalactiae cap*sule types were assigned using GBS-SBG^79^ (git commit 9e53847). *Streptococcus pneumoniae* capsule types were assigned *PneumoKITy*^80^ *v1.0* using default parameters. *Streptococcus pyogenes* genomes were typed using emm-typer v0.2.0 using default parameters (https://github.com/MDU-PHL/emmtyper). *Vibrio parahaemolyticus* capsule types O and K were defined with Kaptive3 using the capsule typing database^81^.

### *Bacillus cereus* group-specific analysis

All sylph-identified *Bacillus cereus sensu lato* (*B. cereus* group) genomes (GTDB genus classification - “Bacillus_A”; n=4,867 genome assemblies) were typed, classified and annotated using BTyper3^82^ v3.4.0 (executed as a module via Bactopia v3.2.0 and Nextflow v25.10.0). Specifically, each assembled genome was supplied as input to BTyper3, (via “bactopia --wf btyper3”, with options “--bt_overlap 0.7” and “--bt_opts --ani_geneflow True”, in addition to the default options), which was used to perform standardized *B. cereus* group-specific taxonomic classification and sequence typing, as well as detection of virulence factors and insecticidal toxin-encoding genes.

### Genome annotation

All assembly FASTA files were annotated using Bakta^83^ v1.9.4 and its full database type v5.1. For consistency and downstream interoperability reasons, the *--keep-contig-headers* option was set; all other parameters were left to default values. To reduce the total amount of genome annotation data (down from 35Tb), a two-step approach was carried out. First, only Bakta JSON result files were kept, as standard output files can be reconstructed using Bakta’s *bakta_io* command. Second, all annotated genome files were then grouped into taxonomic batches as explained above and compressed using xz −9. The resulting compressed annotation data was thereby reduced in size to 1.5TB.

### Protein Structure Prediction of Hypothetical Proteins

Of the 9,319,300,441 proteins in the Bakta annotation output for the 2024-08 release of ATB, we generated protein structures for those that were annotated as hypothetical proteins. After de-duplication of identical sequences, 31,929,327 remained. We then kept only proteins from accessions that passed all quality control checks (as of July 2025), leaving 17,711,165 unique proteins under 3000 amino acids in length. Protein structures were generated using ColabFold^84^ v1.5.5. Multiple sequence alignments were generated with MMSeqs2^85^ v15.6f452 using the uniref30 and environmental (i.e. ColabFoldDB) databases. Protein structure predictions were generated using AMD MI250x GPUs on Setonix at the Pawsey Supercomputing Research Centre using the AlphaFold2^27^ ptm model 1 only, with 3 recycles, no templates or relaxation for maximum throughput.

### APEX predictions

Trained on our in-house peptide and public AMP data, APEX 1.1 is a deep learning-based antimicrobial activity predictor for peptide sequences against *A. baumannii* ATCC 19606*, E. coli* ATCC 11775*, E. coli* AIC221, colistin-resistant *E. coli* AIC222*, K. pneumoniae* ATCC 13883, *P. aeruginosa* PAO1*, P. aeruginosa* PA14*, S. aureus* ATCC 12600, methicillin-resistant *S. aureus* ATCC BAA-1556, vancomycin-resistant *E. faecalis* ATCC 700802, and vancomycin-resistant *E. faecium* ATCC 700221. We used APEX 1.1 to virtually screen all peptides within all bakta-annotated genes within the study. Since APEX has a size limit of 50 amino acids, we provided it with all substrings of length between 8 and 50 of all proteins in the bakta annotations. We used median minimum inhibitory concentrations (MIC) <= 80 μmol L^−1^ to define antimicrobial candidates.

### Striped Smith-Waterman alignment-based similarity Score

We use *SSW(a,b)* to denote the optimal local alignment score of protein sequences *a* and *b* using the Striped Smith–Waterman algorithm^86^ (SSW). The SSW similarity score is defined as

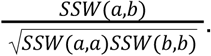

### Peptide sequence space visualization

For the peptide dataset, we computed a pairwise similarity matrix capturing sequence-level similarity among peptides. This matrix was subsequently embedded into a two-dimensional representation using Uniform Manifold Approximation and Projection (UMAP). The resulting low-dimensional embedding provides an approximation of the peptide sequence space, enabling visualization of the global distribution and relationships among peptides.

### Physicochemical properties analysis

The ten physicochemical properties of peptides, including normalized hydrophobic moment, normalized hydrophobicity, net charge, disordered conformation propensity, propensity to aggregation *in vitro*, isoelectric point, linear moment, penetration depth, angle subtended by the hydrophobic residues, and amphiphilicity index, were obtained from the DBAASP server^87^. Note that Eisenberg and Weiss scale^88^ was chosen as the hydrophobicity scale.

### Peptide synthesis and characterization

Peptides were synthesized on an automated peptide synthesizer (Symphony X, Gyros Protein Technologies) by standard 9-fluorenylmethyloxycarbonyl (Fmoc)-based solid-phase peptide synthesis (SPPS) on Fmoc-protected amino acid-Wang resins (100–200 mesh). In addition to preloaded resins, standard Fmoc-protected amino acids were employed for chain elongation. N,N-Dimethylformamide (DMF) was used as the primary solvent throughout synthesis. Stock solutions included: 500 mmol L^−1^ Fmoc-protected amino acids in DMF, a coupling mixture of HBTU (450 mmol L^−1^) and N-methylmorpholine (NMM, 900 mmol L^−1^) in DMF, and 20% (v/v) piperidine in DMF for Fmoc deprotection. After synthesis, peptides were deprotected and cleaved from the resin using a cleavage cocktail of trifluoroacetic (TFA)/triisopropylsilane (TIS)/dithiothreitol (DTT)/water (92.8% v/v, 1.1% v/v, 0.9% w/v, 4.8%, w/w) for 2.5 hours with stirring at room temperature. The resin was removed by vacuum filtration, and the peptide-containing solution was collected. Crude peptides were precipitated with cold diethyl ether and incubated for 20 min at −20 °C, pelleted by centrifugation, and washed once more with cold diethyl ether. The resulting pellets were dissolved in 0.1% (v/v) aqueous formic acid and incubated overnight at −20 °C, followed by lyophilization to obtain dried peptides. For characterization, peptides were dried, reconstituted in 0.1% formic acid, and quantified spectrophotometrically. Peptide separations were performed on a Waters XBridge C_18_ column (4.6 × 50 mm, 3.5 µm, 120 Å) at room temperature using a conventional high-performance liquid chromatography (HPLC) system. Mobile phases were water with 0.1% formic acid (solvent A) and acetonitrile with 0.1% formic acid (solvent B). A linear gradient of 1–95% B over 7 min was applied at 1.5 mL min^−1^. UV detection was monitored at 220 nm. Eluates were analyzed on Waters SQ Detector 2 with electrospray ionization in positive mode. Full scan spectra were collected over m/z 100–2,000. Selected Ion Recording (SIR) was used for targeted peptides. Source conditions were capillary voltage 3.0 kV, cone voltage 25-40 V, source temperature 120 °C, and desolvation temperature 350 °C. Mass spectra were processed with MassLynx software. Observed peptide masses were compared with theoretical values, and quantitative analysis was based on integrated SIR peak areas.

### Bacterial Strains and Growth Conditions

The bacterial panel utilized in this study consisted of the following pathogenic strains: *Acinetobacter baumannii* ATCC 19606; *A. baumannii* ATCC BAA-1605 (resistant to ceftazidime, gentamicin, ticarcillin, piperacillin, aztreonam, cefepime, ciprofloxacin, imipenem, and meropenem); *Escherichia coli* ATCC 11775; *E. coli* AIC221 [MG1655 phnE_2::FRT, polymyxin-sensitive control]; *E. coli* AIC222 [MG1655 pmrA53 phnE_2::FRT, polymyxin-resistant]; *E. coli* ATCC BAA-3170 (resistant to colistin and polymyxin B); *E. coli* K-12 BW25113; *Enterobacter cloacae* ATCC 13047; *Klebsiella pneumoniae* ATCC 13883; *K. pneumoniae* ATCC BAA-2342 (resistant to ertapenem and imipenem); *Pseudomonas aeruginosa* PAO1; *P. aeruginosa* PA14; *P. aeruginosa* ATCC BAA-3197 (resistant to fluoroquinolones, β-lactams, and carbapenems); *Salmonella enterica* ATCC 9150; *S. enterica* subsp. *enterica* Typhimurium ATCC 700720; *Bacillus subtilis* ATCC 23857; *Staphylococcus aureus* ATCC 12600; *S. aureus* ATCC BAA-1556 (methicillin-resistant); *Enterococcus faecalis* ATCC 700802 (vancomycin-resistant); and *Enterococcus faecium* ATCC 700221 (vancomycin-resistant). *P. aeruginosa* strains were propagated on Pseudomonas Isolation Agar, whereas all other species were maintained on Luria-Bertani (LB) agar and broth. For each assay, cultures were initiated from single colonies, incubated overnight at 37 °C, and subsequently diluted 1:100 into fresh medium to obtain cells in mid-logarithmic phase.

### Minimal Inhibitory Concentration (MIC) Determination

MIC values were established using the standard broth microdilution method^89^ in untreated 96-well plates. Test peptides were dissolved in sterile water and prepared as twofold serial dilutions ranging from 1 to 64 μmol L^−1^. Each dilution was combined at a 1:1 ratio with LB broth containing 4 × 10^6^ CFU mL^−1^ of the target bacterial strain. Plates were incubated at 37 °C for 24 h, and the MIC was defined as the lowest peptide concentration that completely inhibited visible bacterial growth. All experiments were conducted independently in triplicate.

### Circular dichroism experiments

The circular dichroism experiments were conducted using a J1500 circular dichroism spectropolarimeter (Jasco) in the Biological Chemistry Resource Center (BCRC) at the University of Pennsylvania. Experiments were performed at 25 °C, the spectra graphed are an average of three accumulations obtained with a quartz cuvette with an optical path length of 1.0 mm, ranging from 260 to 190 nm at a rate of 50 nm min^−1^ and a bandwidth of 0.5 nm. The concentration of all EPs tested was 50 μmol L^−1^, and the measurements were performed in water, a mixture of trifluoroethanol (TFE) and water in a 3:2 ratio, and sodium dodecyl sulfate (SDS) in water at 10 mmol L^−1^, with respective baselines recorded prior to measurement. Spectra were smoothed to minimize background effects. Secondary structure fraction values were calculated using the single spectra analysis tool on the server BeStSel^40^.

### Outer membrane permeabilization assays

N-phenyl-1-napthylamine (NPN) uptake assay was used to evaluate the ability of the peptides to permeabilize the bacterial outer membrane. Inocula of *A. baumannii* ATCC 19606 were grown to an OD at 600 nm of 0.4 mL^−1^, centrifuged (9,391 ’g at 4 °C for 10 min), washed and resuspended in 5 mmol L^−1^ HEPES buffer (pH 7.4) containing 5 mmol L^−1^ glucose. The bacterial solution was added to a white 96-well plate (100 mL per well) together with 4 mL of NPN at 0.5 mmol L^−1^. Consequently, peptides diluted in water were added to each well, and the fluorescence was measured at l_ex_ = 350 nm and l_em_ = 420 nm over time for 45 min. The relative fluorescence was calculated using the untreated control (buffer + bacteria + fluorescent dye) and polymyxin B (positive control) as baselines and the following equation was applied to reflect % of difference between the baselines and the sample:

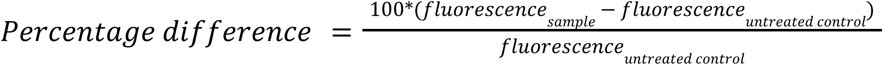

### Cytoplasmic membrane depolarization assays

The cytoplasmic membrane depolarization assay was performed using the membrane potential-sensitive dye 3,3’-dipropylthiadicarbocyanine iodide (DiSC_3_-5). *A. baumannii* ATCC 19606 in the mid-logarithmic phase were washed and resuspended at 0.05 OD mL^−1^ (optical value at 600 nm) in HEPES buffer (pH 7.2) containing 20 mmol L^−1^ glucose and 0.1 mol L^−1^ KCl. DiSC_3_-5 at 20 μmol L^−1^ was added to the bacterial suspension (100 mL per well) for 15 min to stabilize the fluorescence which indicates the incorporation of the dye into the bacterial membrane, and then the peptides were mixed 1:1 with the bacteria to a final concentration corresponding to their MIC values. Membrane depolarization was then followed by reading changes in the fluorescence (l_ex_ = 622 nm, l_em_ = 670 nm) over time for 60 min. The relative fluorescence was calculated using the untreated control (buffer + bacteria + fluorescent dye) and polymyxin B (positive control) as baselines and the following equation was applied to reflect % of difference between the baselines and the sample:

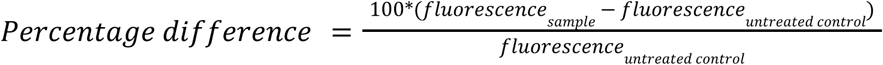

### Cytotoxicity assays

The cells were cultured in high-glucose Dulbecco’s modified Eagle’s medium supplemented with 1% penicillin and streptomycin (antibiotics) and 10% fetal bovine serum and grown at 37 °C in a humidified atmosphere containing 5% CO_2_.

One day prior to the experiment, 100 μL aliquots of human embryonic kidney (HEK293T) cells, at a concentration of 50,000 cells per mL, were seeded into each well of 96-well plates (5,000 cells per well). Following cell attachment, the HEK293T cells were treated with increasing concentrations of peptides (ranging from 8 to 128 μmol L^−1^) and incubated for 24 h. After the exposure period, cytotoxicity was assessed using the 3-(4,5-dimethylthiazol-2-yl)-2,5-diphenyltetrazolium bromide (MTT) assay. Specifically, the MTT reagent was prepared at a concentration of 0.5 mg mL^−1^ in phenol red-free medium and used to replace the peptide-containing supernatants (100 μL per well). The plates were then incubated for 4 h at 37 °C in a humidified atmosphere with 5% CO₂, facilitating the formation of insoluble formazan crystals. These crystals were subsequently dissolved in 0.04 mol L^−1^ hydrochloric acid prepared in anhydrous isopropanol. Absorbance was measured at 570 nm using a spectrophotometer to quantify cell viability. All experiments were conducted in triplicate (three biological replicates).

### Skin abscess infection mouse model

The back of six-week-old female CD-1 mice under anesthesia were shaved and injured with a superficial linear skin abrasion made with a needle. An aliquot of *A. baumannii* ATCC 19606 (1.1×10^6^ CFU mL^−1^; 20 mL) previously grown in LB medium until 0.5 OD mL^−1^ (optical value at 600 nm) and then washed twice with sterile PBS (pH 7.4, 9,391 ’g for 3 min) was added to the scratched area. Peptides diluted in sterile water at their MIC value were administered to the wounded area 1 h post-infection. Two- and four-days post-infection, animals were euthanized, and a uniform excision of the scarified skin was excised, homogenized using a bead beater (25 Hz for 20 min), 10-fold serially diluted, and plated on McConkey agar plates for CFU quantification. The experiments were performed using six mice per group. Mice were single-housed to avoid cross-contamination and maintained under a 12-hour light/dark cycle at 22 °C with humidity controlled at 50%. The skin abscess infection mouse model was revised and approved by the University Laboratory Animal Resources (ULAR) from the University of Pennsylvania (Protocol 806763).

### Reproducibility of experimental assays

All assays were performed in three independent biological replicates as indicated in each figure legend and in the Experimental Models and Methods details sections. The values obtained for hemolytic activity were estimated by non-linear regression based on the screen of peptides in a gradient of concentrations and represent the hemolytic concentration values needed to lyse and kill 50% of the cells present in the experiment. In the skin abscess and thigh infection mouse models, we used six mice per group following established protocols approved by the University Laboratory of Animal Resources (ULAR) of the University of Pennsylvania.

### Quantification and statistical analysis

In the mouse experiments, all the raw data were log_10_ transformed and the statistical significance was determined using one-way ANOVA followed by Dunnett’s test. All the p values are shown for each of the groups, all groups were compared to the untreated control group. All calculation and statistical analyses of the experimental data were conducted using GraphPad Prism v.10.4. Statistical significance between different groups was calculated using the tests indicated in each figure legend. No statistical methods were used to predetermine sample size.

## Code Availability

All code used is available at https://github.com/AllTheBacteria/AllTheBacteria/. The assembly pipelines for release 0.2 and 2024-08 are at https://github.com/leoisl/bacterial_assembly_pipeline and https://github.com/AllTheBacteria/AllTheBacteria/tree/main/reproducibility/All-samples/assembly.

## Data availability

A website linking to all resources is available at https://allthebacteria.org/. Documentation for AllTheBacteria is available at https://allthebacteria.readthedocs.io/en/latest/. All data for AllTheBacteria are hosted on the Open Science Framework (OSF) here: https://osf.io/xv7q9/. Genome annotations in an Apache Parquet dataset (containing gene and intergenic sequences, locations and genomic context for all samples) are available on HuggingFace here: https://huggingface.co/datasets/AllTheBacteria/ATB, with a deduplicated version here https://huggingface.co/datasets/AllTheBacteria/ATB-representative-ids. In addition, individual assembly FASTA files for each sample are available on AWS with S3 URI of the form s3://allthebacteria-assemblies/<SAMPLE_ID>.fa.gz (for example s3://allthebacteria-assemblies/SAMD00000344.fa.gz). Assemblies are also available through the ENA; in their internal terminology these are called third-party annotations (TPAs) with ERZ prefixes. A LexicMap index of the full dataset is also available on AWS (see https://allthebacteria.org/docs/ for details). We provide a metadata browser here https://allthebacteria.org/browse.

We also provide a command-line utility (available at https://github.com/AllTheBacteria/atb-cli) which allows rapid access to summary data, filtering by metadata, AMR gene content and MLST type and finding genomes similar to a user-provided genome.

## Author contributions

Abbreviations refer to author initials. To remove ambiguity: VigR refers to Vignesh Ramnath, and VisR to Vishnu Raghuram. ThS refers to Theo Sanderson, and TiS to Timo Saratto. MH refers to Martin Hunt, and MBH to Michael Hall.

Conceptualisation: MH, LL, JAL, ZI.

Data curation: MH, LL, DA, GB, JH, OS, SMA, STH, NM, MW, AW, WS

Formal Analysis: MH, DA, GB, JH, OS, SMA, JAL, TS, MC, JB, VigR, VisR, MAC, LMC, MJR, STH, JT, JvW, OX, NM, RP3, MW, AW, N-FA, FB, BJ, WM, AMM, RMW, ThS, LPS, H-SL, FW, JAL

Funding acquisition: MBH, AF, NT, CdlF-N, JAL, ZI

Investigation: MH, MDTT, LL, MLA, FB, BJ, WM, FW, JAL, ZI

Methodology: MH, MDTT, JT, JvW, OX, TLV, FW, GT-H, WS, CdlF-N, JAL, ZI

Project Administration: CdlF-N, JAL, ZI

Resources: GB, MBH, MW, AW, AF, ThS, TiS, NT, CdlF-N, JAL, ZI

Software: MH, LL, OS, TS, JT, JvW, MW, AW, AF, TiS, TLV, WS, CdlF-N, JAL, ZI

Supervision: TS, AF, NT, CdlF-N, JAL, ZI

Validation: MH, MDTT, TS

Visualisation: MH, MDTT, NM, RMW, LPS, JAL

Writing - original draft: MH, MDTT, JH, WM, RMW, LPS, CdlF-N, JAL, ZI

Writing - review and editing: all authors.

## Acknowledgements

We would like to thank Karel Břinda for help with running MiniPhy. We would like to thank Fabien Voisin and Sarah Beecroft for their assistance in operating ColabFold at scale at Phoenix and Pawsey respectively, with extra acknowledgement to Sarah for containerising ColabFold for use on Setonix’s AMD GPUs. We thank Dr. Mark Goulian for kindly donating the following strains: *Escherichia coli* AIC221 [*Escherichia coli* MG1655 phnE_2::FRT (control strain for AIC222)] and *Escherichia coli* AIC222 [*Escherichia coli* MG1655 pmrA53 phnE_2::FRT (polymyxin resistant)]. We thank the de la Fuente lab for insightful discussions. We thank UKHSA for the rapid sequencing and public release of genomes from the 2026 Kent meningitis outbreak.

We are grateful to Amazon AWS for an AWS Open Data Sponsorship Award which funds hosting of assembly data and a LexicMap index. This work was supported with the assistance of resources and services from Phoenix HPC at the University of Adelaide and Pawsey Supercomputing Research Centre, which is supported by the Australian Government. MH was supported by Sanger Institute core funding from the Wellcome Trust [220540/Z/20/A], and by the National Institute for Health Research (NIHR) Health Protection Research Unit in Healthcare Associated Infections and Antimicrobial Resistance at Oxford University in partnership with the United Kingdom Health Security Agency (UKHSA) (NIHR200915), and also supported by the NIHR Biomedical Research Centre, Oxford. The views expressed in this publication are those of the authors and not necessarily those of the NHS, the National Institute for Health Research, the Department of Health or the UKHSA. JH was supported by an Emerging Leadership Fellowship from the National Health and Medical Research Council (NHMRC) of Australia (2034741). This work was supported by Monash eResearch capabilities, including the high performance computer M3 and Research Data Storage. OS and this work was supported by the German Network for Bioinformatics Infrastructure (de.NBI) and the de.NBI cloud hosted by the BiGi Service Center within de.NBI and ELIXIR-DE (W-de.NBI-010). MLA was supported by Coordenação de Aperfeiçoamento de Pessoal de Nível Superior - Brasil (CAPES) - Finance Code 001. LMC was supported by SciLifeLab & Wallenberg Data Driven Life Science Program (grant KAW 2020.0239); Swedish Research Council (VR; grant 2023-05212). MW, AW and AF were supported by Wellcome Discovery award 226602/Z/22/Z LifeArc/CF Trust Innovation Hub THUB01. N-FA was supported by Quadram Institute Bioscience BBSRC funded Core Capability Grant (project number BB/CCG1860/1). AMM was supported by the Center for Microbial Medicine at CHOP and the National Institutes of Health grants U19AI174998 and 1R01AI185544-01A1. RMW supported by a UK Research and Innovation Future Leaders Fellowship (UKRI2317). TS was supported by the Wellcome Career Development Award 306920/Z/23/Z. LPS acknowledges an ERC Starting Grant (PLEIADES 101219784). CdlF-N holds a Presidential Professorship at the University of Pennsylvania; research in this publication was supported by the National Institute of General Medical Sciences of the National Institutes of Health under award number R35GM138201 and the Defense Threat Reduction Agency (DTRA; HDTRA1-21-1-0014). WM was supported by the Wellcome Early-Career Award 319534/Z/24/Z. GT-H was supported by the Australian Research Council (DE240100316) and the National Health and Medical Research Council (GNT2025515).

## Materials and Correspondence

For correspondence related to access to the AllTheBacteria data or bioinformatics, please contact. For correspondence relating to the antimicrobial peptide discovery and materials therein, contact.

