## Supplementary Material for "AllTheBacteria: a community resource empowers biology and discovers novel peptide antibiotics"

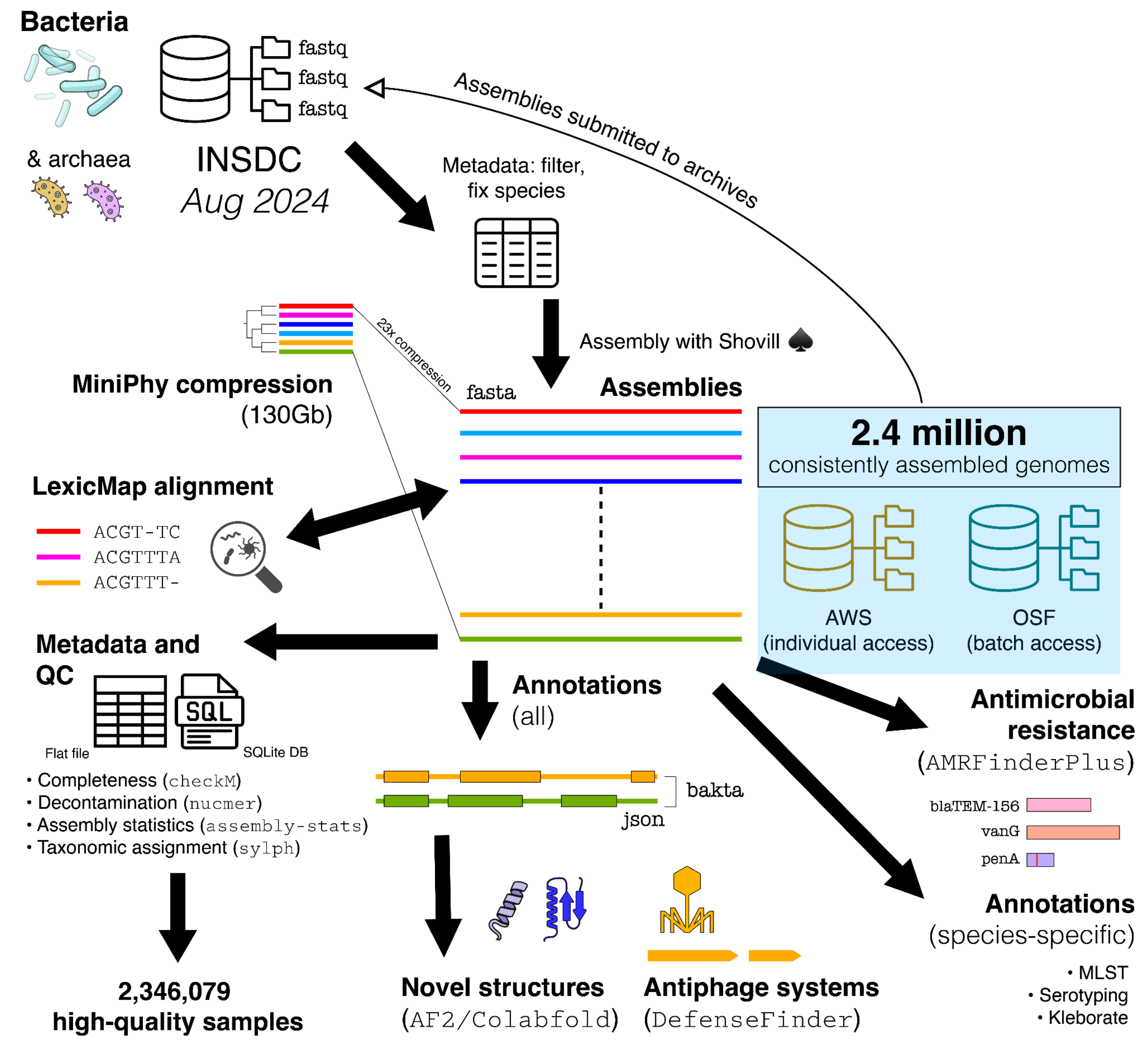


**Supplementary Figure 1. AllTheBacteria workflow and community architecture.** Public bacterial and archaeal short-read sequencing data (fastq files) from the International Nucleotide Sequence Database Collaboration are imported, assembled and subjected to standardized quality control, including independent species verification and contamination assessment. High-quality assemblies are annotated, compressed with MiniPhy, made searchable with LexicMap and distributed through cloud and batch-access routes. Downstream analyses include antimicrobial-resistance profiling, antiphage-defence detection, protein-structure prediction for novel proteins and community-contributed species-specific annotations.

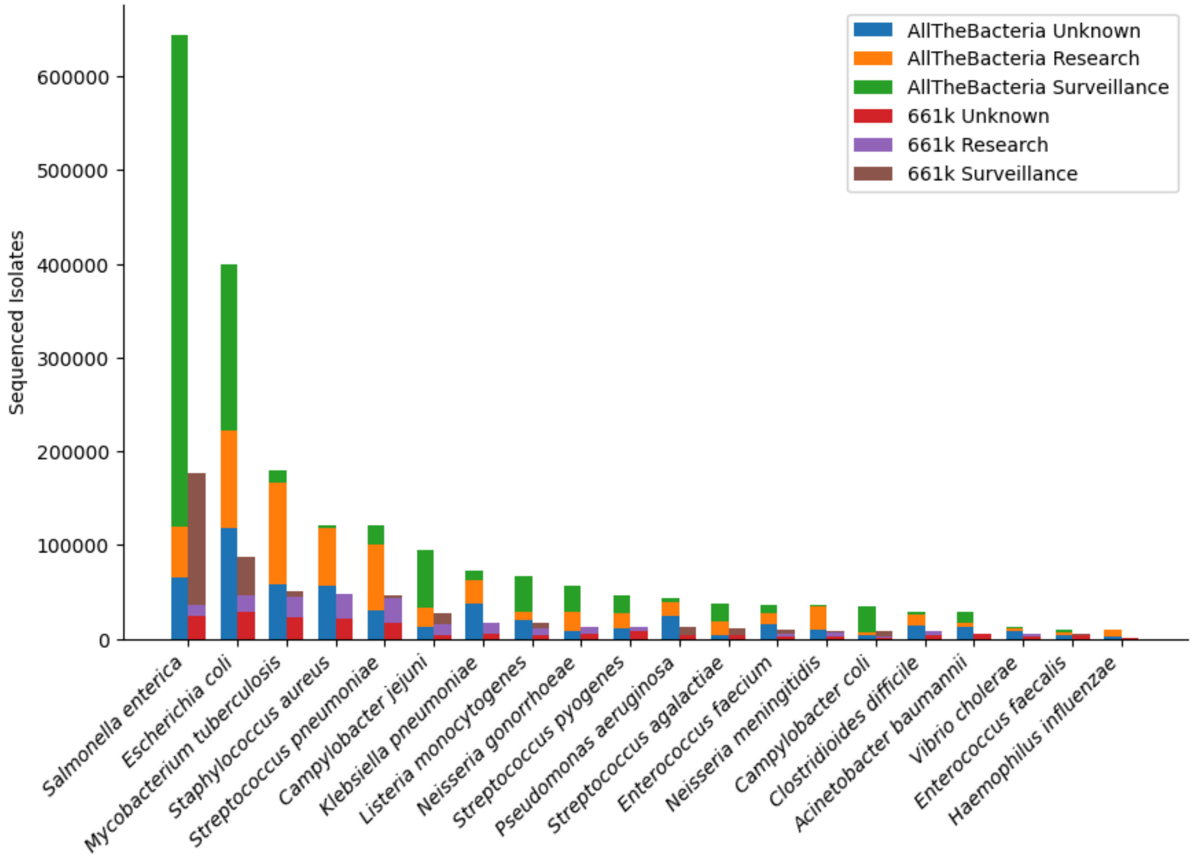


**Supplementary Figure 2**. Sampling purpose across AllTheBacteria and the 661k reference dataset. The bar plot compares the most frequently sampled species and partitions isolates by study purpose, showing the expanded scale and surveillance contribution of AllTheBacteria relative to the earlier 661,000-sample dataset.

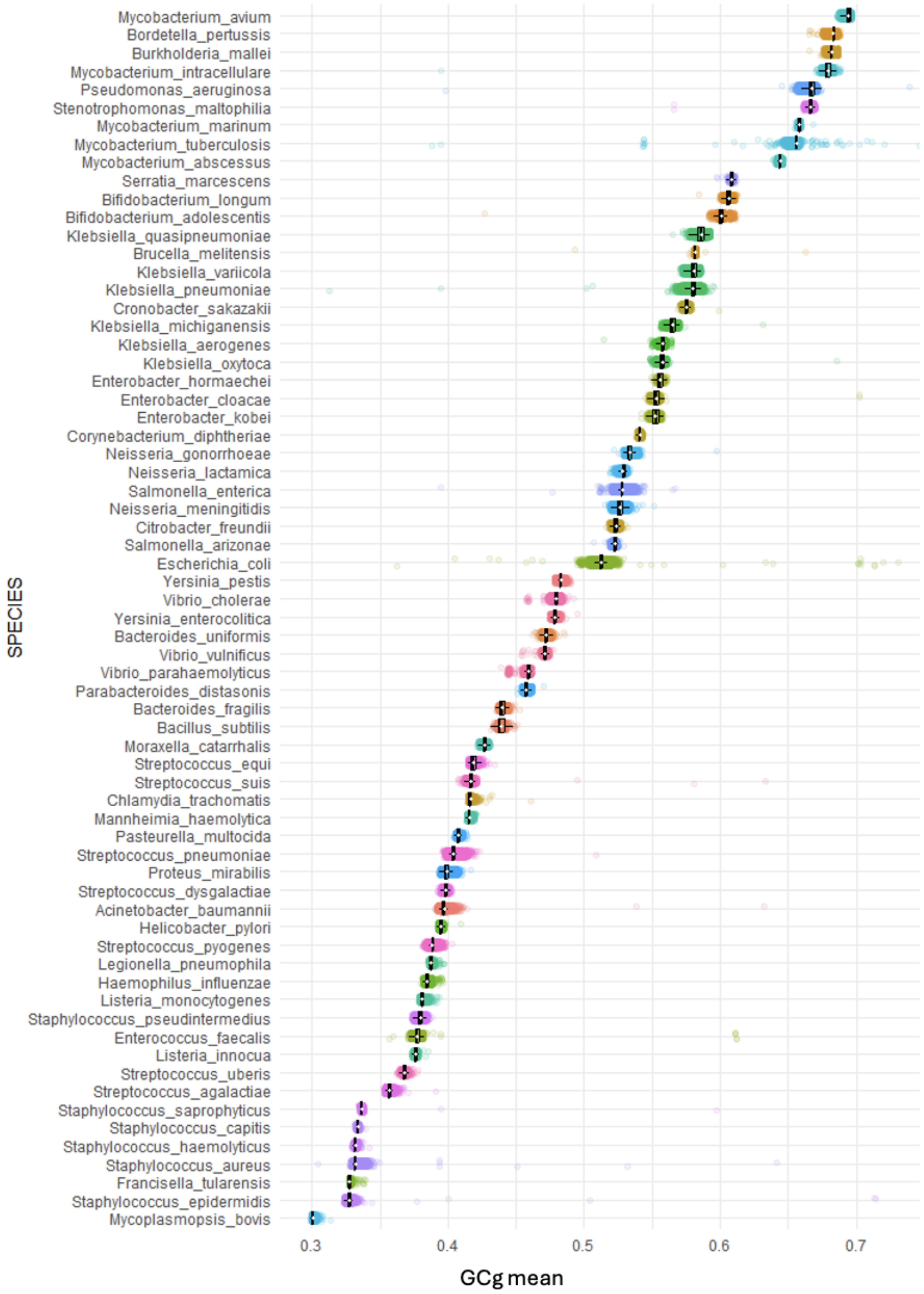


**Supplementary Figure 3.** Per-species distributions of per-predicted gene GC mean (GCg-mean). Each dot represents the GCg-mean of a genome for predicted genes (ORFs) >300bp, ordered from low to high GCg-mean.

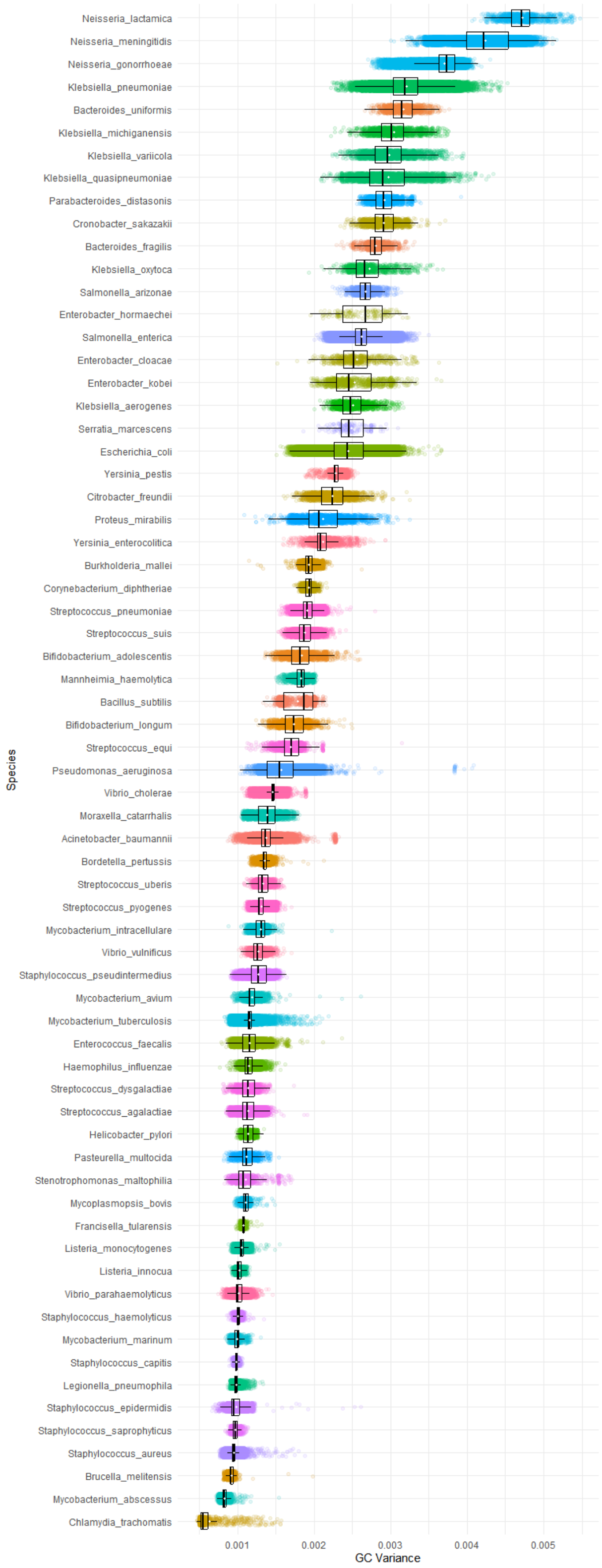


**Supplementary Figure 4.** Per-species distributions of per-gene %GC *variance (GCg-variance)* for each genome for predicted genes >300bp, ordered from highest to lowest variance. Each row shows the distribution of per-assembly variance as a boxplot over jittered points.

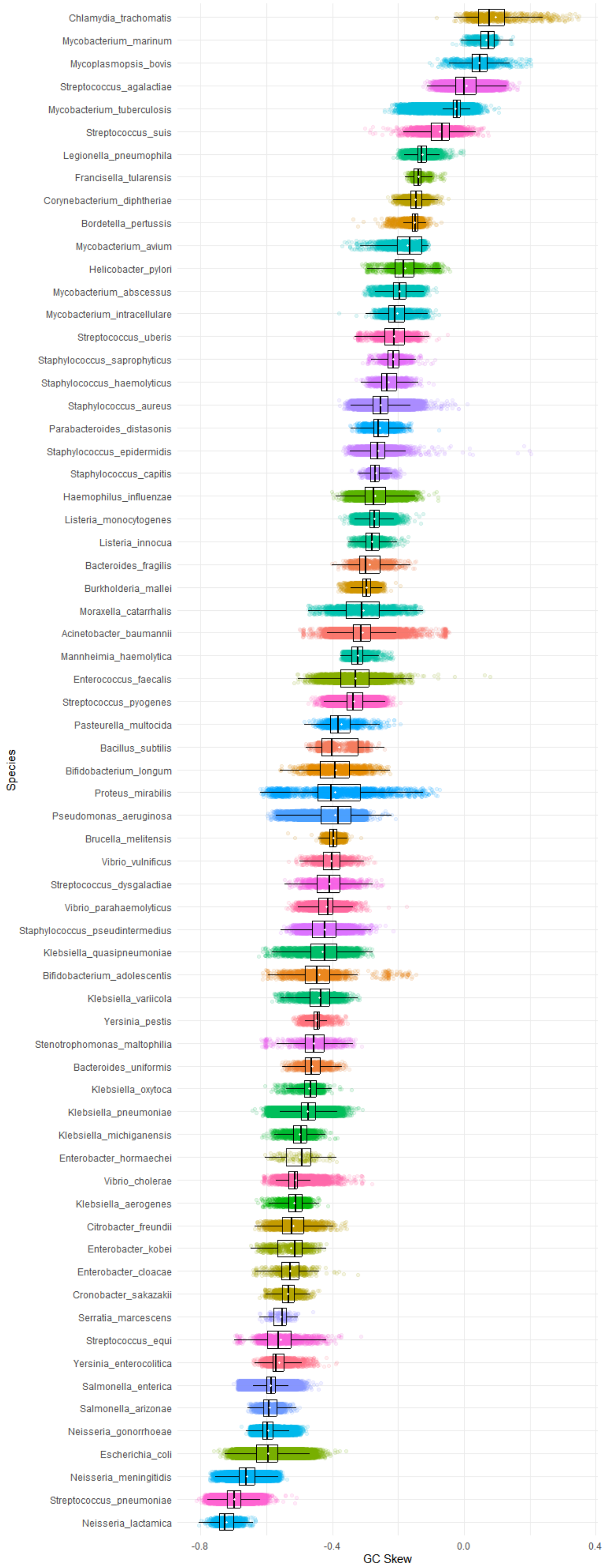


**Supplementary Figure 5.** Per-species distributions of per-gene %GC *skew (GCg-skew)* for each genome for predicted genes >300bpwith each species ordered from most positive (right skewed) to most negative (left skewed). This should not be confused with strand-specific GC skew.

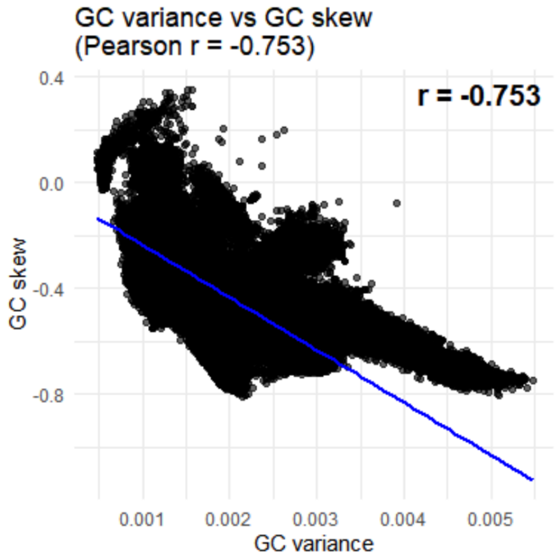


**Supplementary Figure 6.** Correlation between genome assembly level statistics of per-gene GC skew (GCg-skew) for each assembly and per-gene GC variance (GCg-variance) for each assembly.

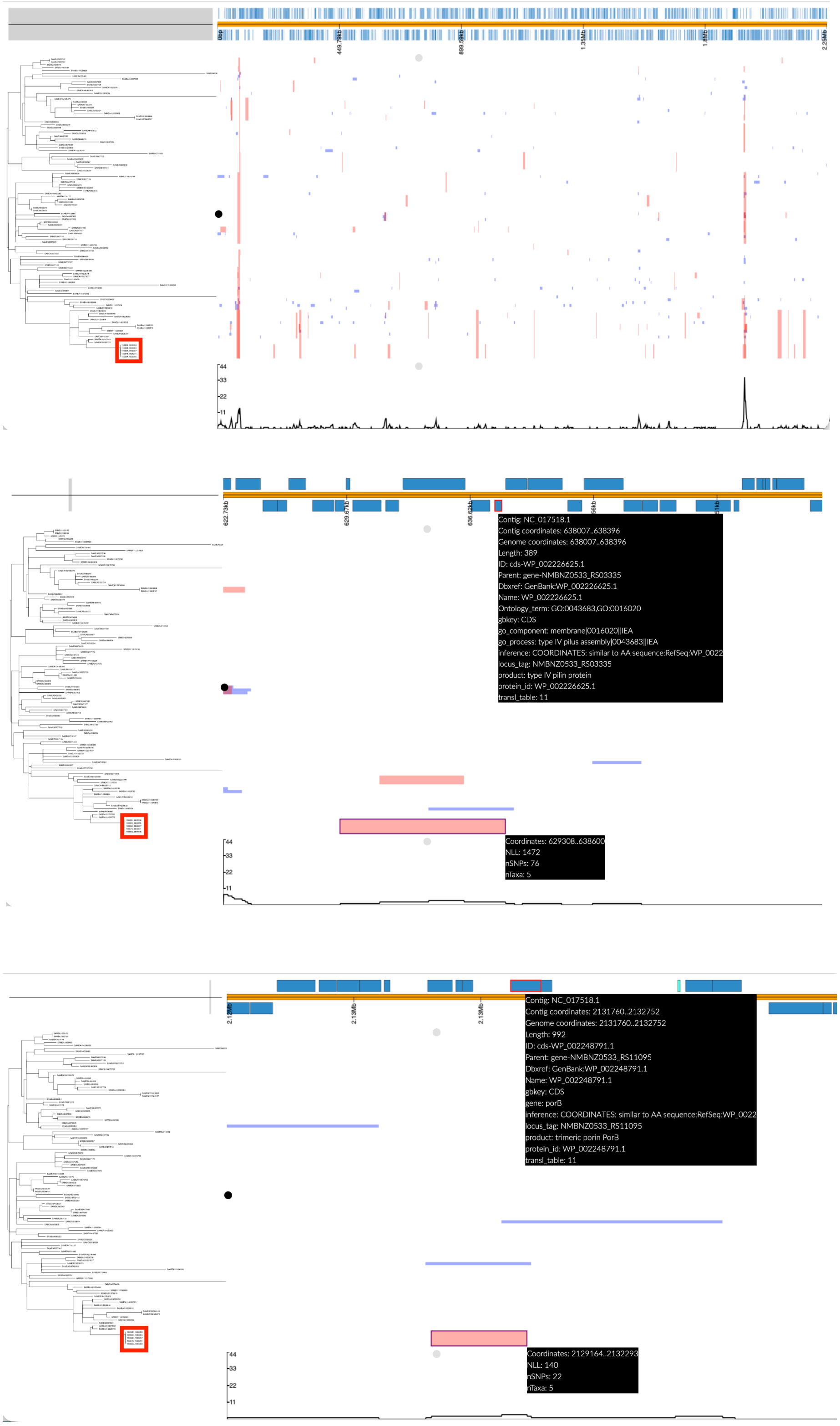


**Supplementary Figure 7. Recombination analysis of the UK 2026 meningitis outbreak.** In each subfigure: phylogenetic tree in the left panel with outbreak genomes in red box; top panel shows genes annotated in reference; central panel shows detected recombination events (red blocks are events at internal nodes, blue at terminal branches); bottom panel shows density of recombination. Screenshots from phandango (https://jameshadfield.github.io/phandango/#/). Top subfigure: recombination detected in outbreak when provided with context of neighbour genomes from AllTheBacteria. Nine recombination events private to the outbreak clade were detected. Middle subfigure: zoom to the recombination event overlapping pilus at 638kb (with annotation highlighted in top panel). Bottom subfigure: zoom to the recombination event overlapping *porB* at 2.13Mb (with annotation highlighted in top panel).

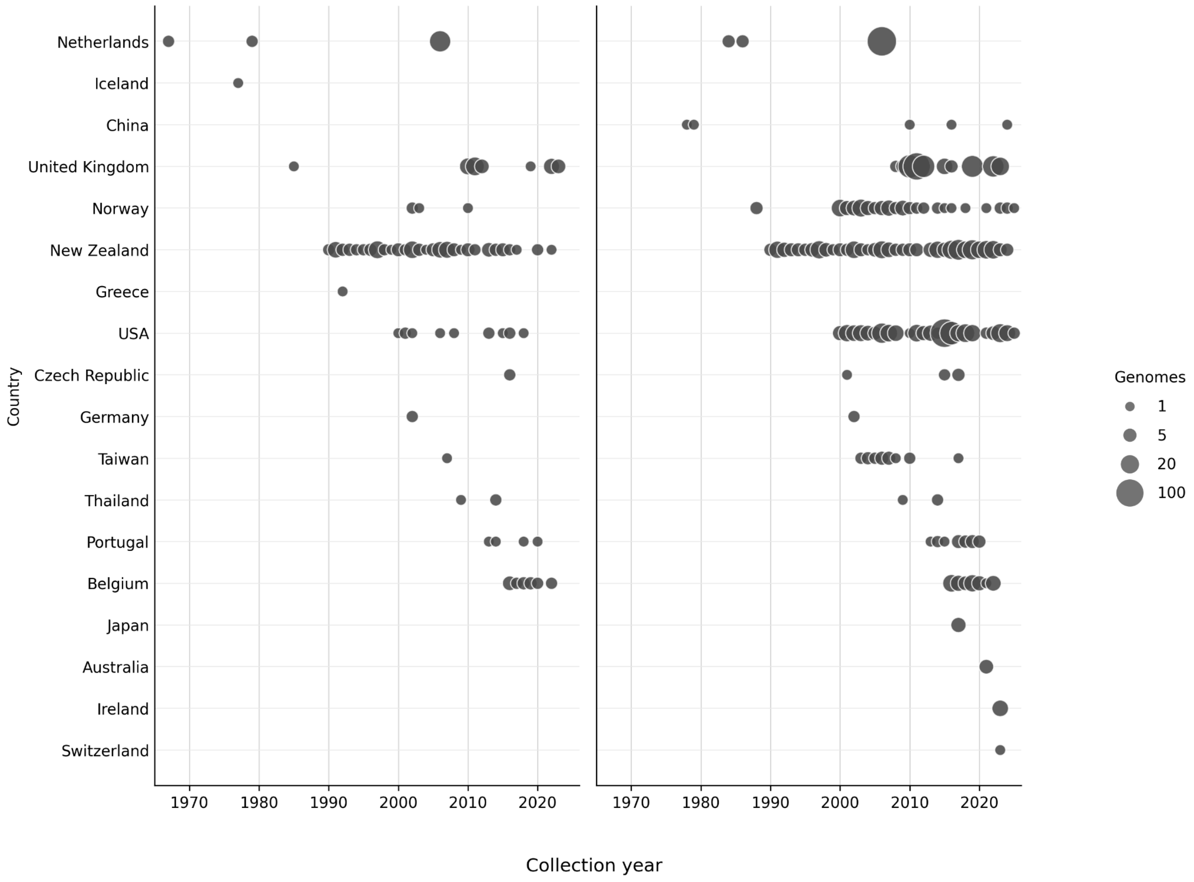


**Supplementary Figure 8. Global and temporal distribution of meningococcal genomes with perfect nucleotide matches to PorB porin and type IV pilin coding sequences.** Country and collection year distribution of *Neisseria meningitidis* genomes with perfect nucleotide matches to the coding sequences encoding type IV pilin WP_002226625.1 (left plot) and PorB porin WP_002248791.1 (right plot). Only genomes that pass AllTheBacteria QC filters are included. Points represent country-year bins and are scaled by the number of genomes in each bin. Collection dates were binned to year; date ranges were assigned to the earliest year, and sub-national locations were collapsed to country. The type IV pilin coding sequence was detected in 239 genomes from 13 countries between 1967 and 2023, and the PorB porin coding sequence was detected in 1192 genomes from 16 countries between 1978 and 2025. These distributions indicate that both antigen-associated coding sequences were present in meningococcal genomes decades before the outbreak period and were geographically widespread.

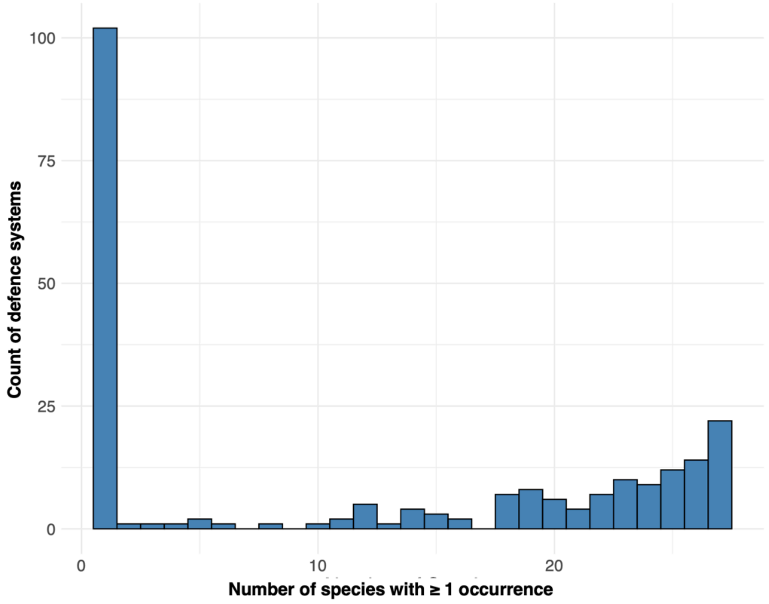


**Supplementary Figure 9. Distribution of antiphage-defence systems across species.** For each defence-system family, we counted the number of species in which the system was detected at least once and plotted the resulting distribution. Most systems were rare: 102 were detected in a single species, including 99 found only in *E. coli*. Nevertheless, 84 systems were detected in more than 20 species, highlighting a core set of broadly distributed defence functions across Enterobacterales.

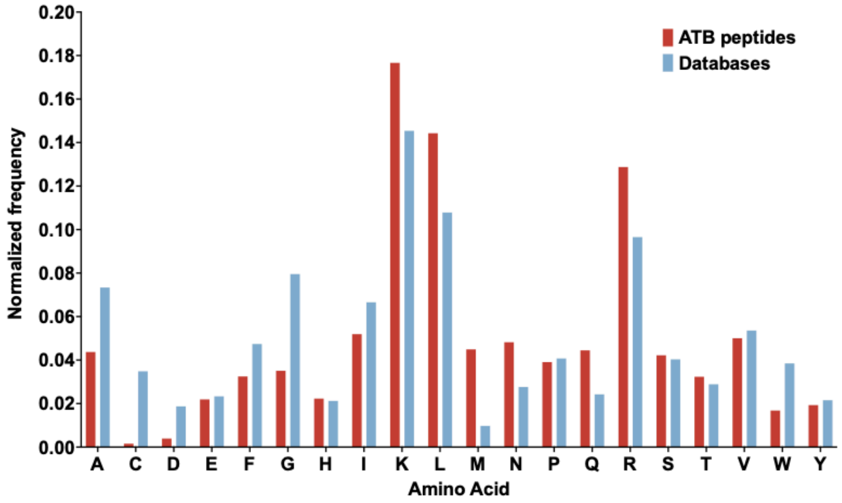


**Supplementary Figure 10. Amino acid composition of APEX-predicted AllTheBacteria peptides compared with known antimicrobial peptides from DBAASP, APD3 and DRAMP 3.0.**

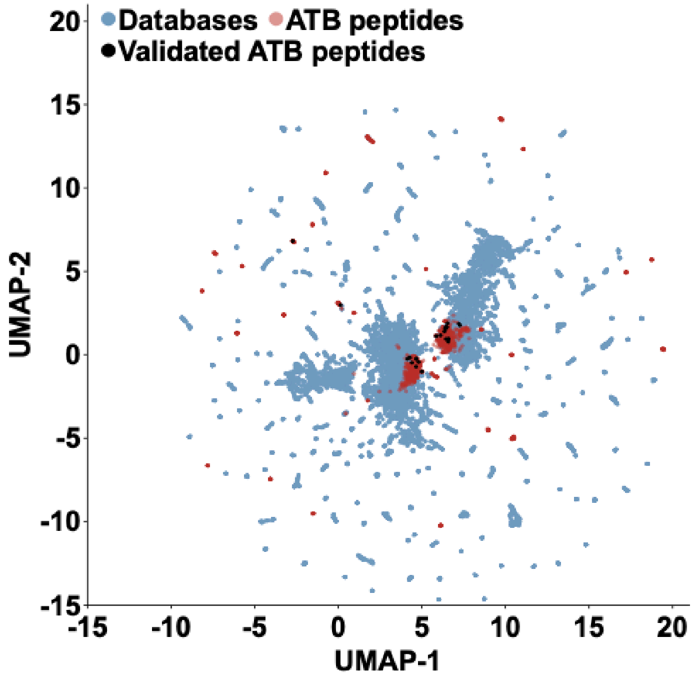


**Supplementary Figure 11. Sequence space exploration of AllTheBacteria-derived peptides and database antimicrobial peptides.**

The graph illustrates a bidimensional sequence space visualization of peptide sequences found in DBAASP and antimicrobial EPs discovered by APEX in ATB. Sequence alignment was used to generate a similarity matrix for all peptide sequences in databases and the ATB-peptides predicted by APEX. Each row in the matrix represents a feature representation of a peptide based on its amino acid composition. Uniform Manifold Approximation and Projection (UMAP) was applied to reduce the feature representation to two dimensions for visualization.

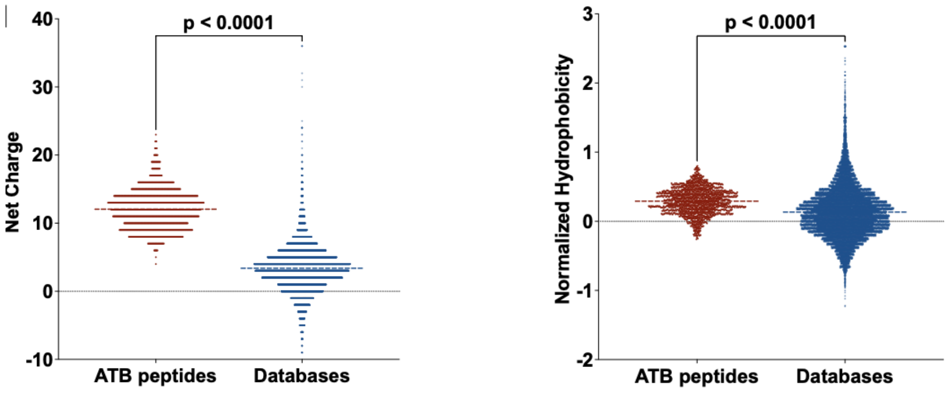


**Supplementary Figure 12. Net charge and hydrophobicity distributions of predicted AllTheBacteria peptides compared with database antimicrobial peptides.**

Distributions are shown for two physicochemical properties associated with antimicrobial activity: (left) net charge, and (right) and normalized hydrophobicity. Net charge influences the initial electrostatic interactions between the peptide and negatively charged bacterial membranes, while hydrophobicity affects interactions with lipids in the membrane bilayers.

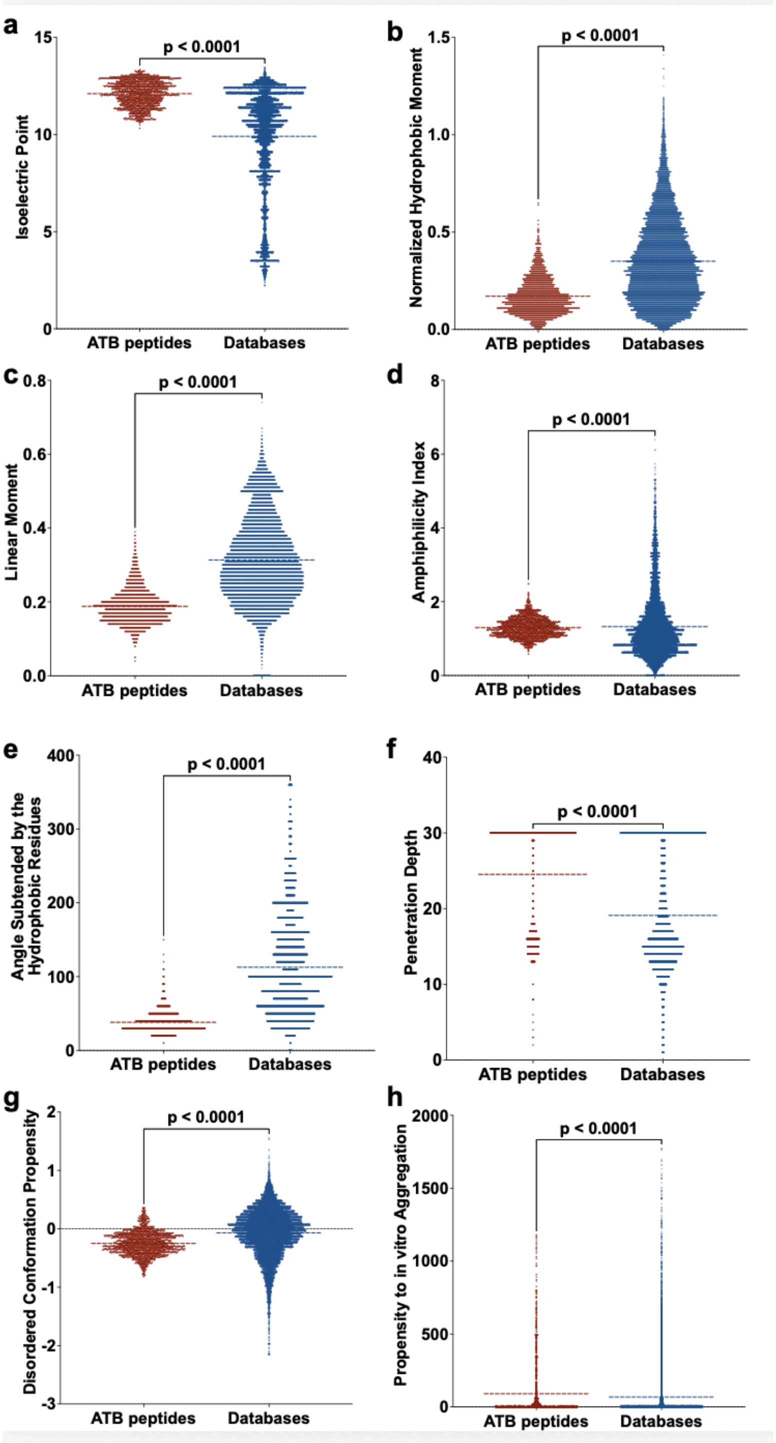


**Supplementary Figure 13**. **Additional physicochemical features of AllTheBacteria peptides compared with database antimicrobial peptides.** **(a)** Isoelectric point, **(b)** normalized hydrophobic moment, and **(c)** linear moment, reflecting the amphipathicity of the molecules, which directly influences their interactions with bacterial membranes. **(d)** Amphiphilicity index, **(e)** angle subtended by the hydrophobic residues, **(f)** penetration depth, and **(g)** disordered conformation propensity, both of which are closely correlated with the mechanism of action, specifically how peptides interact with membrane lipids to exert antimicrobial activity. **(h)** Propensity to aggregate *in vitro*, correlated with the supramolecular arrangement of the molecules and potential toxicity. Statistical significance was determined using two-tailed t-tests followed by the Mann-Whitney test; p values are shown in the graph. The solid line within each box represents the mean value for each group.

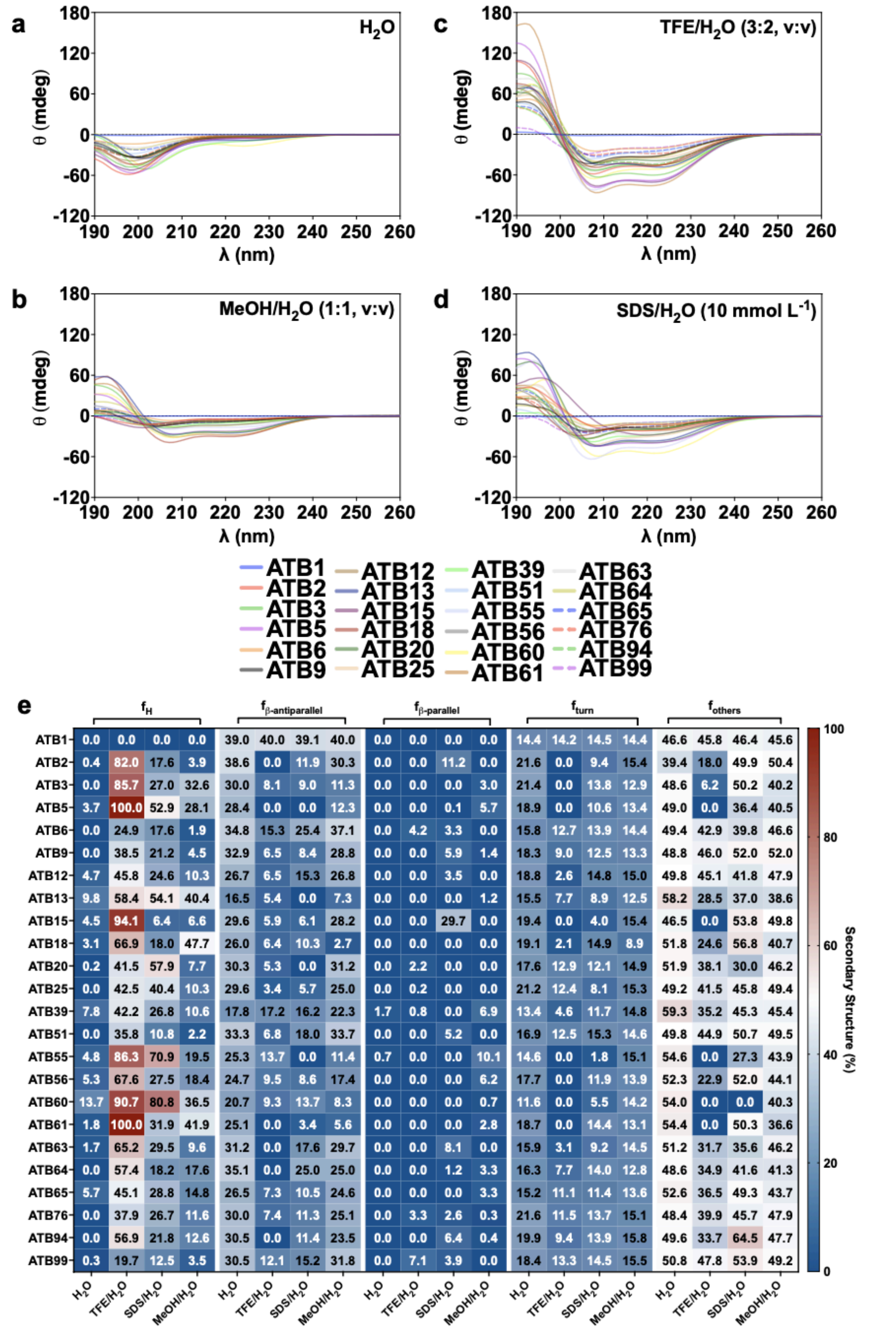


**Supplementary Figure 14. Circular dichroism spectra of ATB peptides.** Circular dichroism experiments were conducted with ATB peptides using a J-1500 Jasco circular dichroism spectrophotometer. The spectra were recorded in four different media: **(a)** water, **(b)** 50% methanol in water, **(c)** 60% trifluoroethanol in water, and **(d)** sodium dodecyl sulfate (SDS) in water (10 mmol $L^{-1}$), after three accumulations at 25 °C, using a 1mm path length quartz cell, between 260 and 190 nm at 50 nm $min^{-1}$, with a bandwidth of 0.5 nm. The concentration of all peptides tested was 50 mmol $L^{-1}$. **(b)** Heatmap with the percentage of secondary structure found for each peptide in the four different solvents. Secondary structure fraction was calculated using the BeStSel server.

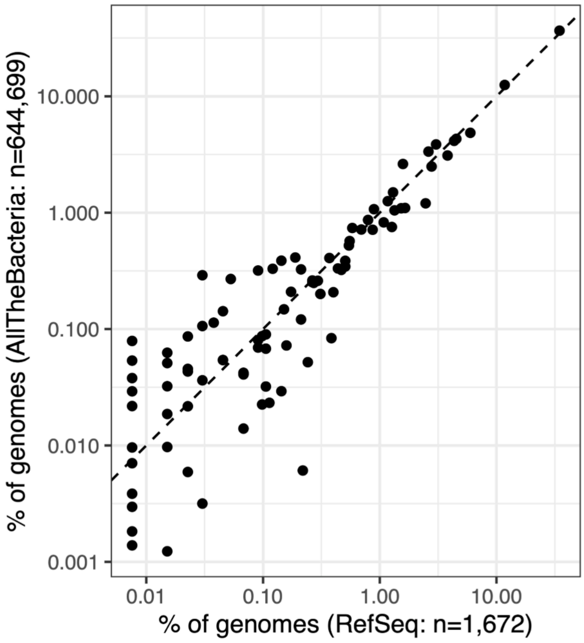


**Supplementary Figure 15. Defence-system detection is consistent between AllTheBacteria assemblies and complete RefSeq *Salmonella enterica* genomes.**
The plot compares the prevalence of defence-system families detected by DefenseFinder in *S. enterica* across the two datasets. Only systems detected at least once in both datasets are shown (91 of 119 systems detected in AllTheBacteria assemblies); 28 additional systems were detected in AllTheBacteria).

### Supplementary Note 1. Predicted gene (ORF) level %GC content statistics aid genome QA/QC and species/replicon classification, and could be a proxy for microbial clonality and ecological isolation/specialization

Using the ATB dataset consisting of assemblies processed through one uniform annotation pipeline, we calculated the %GC of every predicted coding gene (ORF) > 300 bp, in any genome assembly where there were >400 genomes/species. We then calculated three summary statistics: the mean and variance of this per-gene level %GC for each genome assembly, as well as the estimated skewness with the Pearson median skewness formula (skew ≈ 3 × (mean − median) / SD; this should not be confused with strand-specific GC skew). In addition, species-level values were calculated of the mean across that species' assemblies. Previously we had identified trends in such per-gene %GC statistics based on genes using a very limited dataset (https://www.pathogenomics.sfu.ca/islandpath/update/IPindex.pl?stddev), however the ATB dataset's annotation consistency now provides an opportunity to calculate this on a large scale, using consistent criteria not confounded by differing assembly approaches or gene-calling methods. This analysis highlights how consistent methods can both aid QA/QC, and reveal insights into genome evolution and potential clonality of a species, detected from a single genome.

**QA/QC, species classification, replicon identification utility:**
Per-gene %GC analysis for ATB genome assemblies was useful in exposing potential data-quality and/or species classification anomalies. When we plotted the per-gene GC mean for each genome (GCg-mean) of a given species, most species had a range of GCg-mean distributions that are centered around the mean of that species. Outliers in the GCg-mean distribution can indicate potential species misclassification. For example, assembly SAMN36357135 was annotated as *Klebsiella pneumoniae*, with a GCg-mean of 39%. However, the mean for *K. pneumoniae* in the ATB data is 57% (Suppl. Fig. 3). In the ENA, this assembly was annotated as *Acinetobacter sp.* 2019-01-40-01, which is more consistent with the GCg-mean of 39% observed in ATB for *Acinetobacter baumannii*. A simple GCg-mean calculation allowed such discrepancies to be rapidly detected.

In addition there were cases where the gene-based %GC for genomes for a species were more bimodal when plotted (Supp. Fig. 3). Some are explainable, for example,

*Vibrio* species have bimodal distributions for GCg-mean, consistent with their two chromosomes/replicons genomes. In other bimodal cases, species may be more suitably subclassified.

Generally, for newly characterized single genomes, such analysis can quickly support classification and/or provide further evidence for multiple replicons.

**Aiding identification of genome/species clonality and ecological isolation:** When we calculated per-gene %GC *variance* for each genome (GCg-variance) we observed marked differences in variance across genomes (Supp. Fig. 4). Notably, genomes with lowest GCg-variance were *Chlamydia trachomatis*, an obligate intracellular pathogen that is relatively ecologically isolated versus other bacteria. Conversely, *Neisseria* species had the highest GCg-variance. *Neisseria* are noted for their natural competence for DNA uptake. While this uptake is facilitated by a species-associated uptake sequence, their high GCg-variance suggests that their frequent horizontal transfer of DNA is impacting their genomes on a broad scale. A similar observation was noted for per-gene %GC *skew* for each genome (GCg-skew; Suppl. Fig. 5). The most right-skewed (positive) species is *Chlamydia trachomatis* while the most strongly left-skewed (negative) species are the naturally transformable *Neisseria*. This per-gene variance and skew for an individual genome is negatively correlated with each other (Suppl. Fig. 6), such that lower variance in per-gene GC for a genome is associated with higher skew.

These results suggest that calculating per-gene GC variance and skew for an individual genome of a novel species, may provide a surprisingly simple first-pass, fully sequence-based estimation of if that species or lineage is particularly clonal, or ecologically isolated/specialized, verses more subject to frequent horizontal gene transfer/DNA uptake.

In addition, genomes subject to less horizontal DNA transfer, like *Chlamydia*, are expected to converge on a characteristic %GC distribution, consistent with its lower per-gene %GC variance, however, the distribution skew is high. We hypothesize that for more ecologically isolated species they will converge on an optimal %GC, in absence of horizontally acquired DNA, however a constrained subset of genes will remain relatively recalcitrant to this optimal GC content, skewing the distribution.

Collectively, per-gene %GC, calculated for a genome assembly, can serve as a lightweight, automatable screen for aiding genome assembly, replicon and taxonomic assignment QA/QC. In addition, even the simple per-gene %GC analysis of one novel bacterial genome may provide some insight into clonality/ecological isolation/specialization of a species. Further analyses, performed consistently over this ATB dataset, could also help address further questions regarding evolution of %GC.

References: https://pmc.ncbi.nlm.nih.gov/articles/PMC4450053/, https://pubmed.ncbi.nlm.nih.gov/16200051/, https://link.springer.com/article/10.1186/1471-2164-11-464.

### Supplementary Note 2. Example application - meningitis outbreak

Analyses were performed using the AllTheBacteria command line tool 'atb' version 0.17.1, and AllTheBacteria releases up to and including 2025-05. Closely related genomes to the 5 outbreak genomes (PubMLST IDs 190673_1926231, 190682_1930357, 190683_1930355, 190684_1930356, 190685_1930358) were identified using 'atb sketch query --format tsv --knn 500', retaining the best 500 matches. Because multiple hits may originate from the same genome, unique genome accessions were extracted from the matches, yielding 100 distinct samples. Metadata for these samples were retrieved using 'atb query', and genome assemblies were downloaded using 'atb download'.

A core-genome alignment of the 5 outbreak genomes and the 100 nearest genomes was generated using Gubbins version 3.4.3. First, a whole-genome alignment was produced with generate_ska_alignment.py using GCF_000191525.1 as the reference genome. Recombination filtering and phylogenetic inference were then performed with run_gubbins.py using a filter percentage of 30%.

The nucleotide sequences corresponding to two antigen-associated proteins of interest were extracted from the reference genome using SeqKit: a type IV pilin protein, WP_002226625.1, at 638,007-638,396, and the PorB porin, WP_002248791.1, at 2,131,760-2,132,752. These were chosen based on their known role as antigens (https://doi.org/10.1016/j.vaccine.2022.05.032, https://link.springer.com/protocol/10.1007/978-1-61779-346-2_1). Nucleotide sequence similarity searches were performed with LexicMap version 0.9.0 using default parameters against a database constructed from AllTheBacteria releases up to 2025-05. The nucleotide sequences corresponding to WP_002226625.1 and WP_002248791.1 returned 104,011 and 169,481 matches, respectively, with 2,020 and 4,636 samples containing a perfect match. Metadata associated with matching samples were retrieved using atb query --columns sample_accession,hq_filter,sylph_species,country,collection_date, allowing extraction of sample accession, quality-control status, species assignment, country of origin, and collection date.

### Supplementary Note 3. Distinctive physicochemical features of antimicrobial peptides discovered with APEX

**Comparison with known antimicrobial peptide databases**

We compared the amino acid composition of APEX-predicted AllTheBacteria peptides with experimentally annotated antimicrobial peptides from DRAMP, APD3 and DBAASP. The AllTheBacteria peptides were strongly enriched in basic residues, especially lysine and arginine, supporting electrostatic engagement with negatively charged bacterial envelopes. Many sequences also contained hydrophobic and aromatic residues, including leucine, isoleucine and tryptophan, suggesting the capacity to form amphiphilic motifs that interact with, insert into or disrupt bacterial membranes.

To position these sequences within the broader antimicrobial peptide landscape, we embedded both APEX-predicted AllTheBacteria peptides and database peptides into a UMAP sequence-similarity map (Supplementary Fig. 5). Known antimicrobial peptides formed recognizable clusters corresponding to recurrent sequence families, whereas AllTheBacteria-derived candidates occupied both overlapping regions and sparsely populated areas of the map. This pattern suggests that encrypted bacterial peptides extend the known antimicrobial sequence landscape and provide access to scaffolds not well represented in current experimental databases.

**Physicochemical features of ATB peptides**

To characterise the physicochemical space occupied by APEX-predicted ATB peptides, we used the DBAASP "properties" tools to calculate a panel of descriptors commonly employed in antimicrobial peptide analysis and compared their distributions with those of reference AMPs curated in DRAMP, APD3 and DBAASP (see Supp. Fig 3). Across the full APEX-prioritised ATB set, the most striking difference relative to database AMPs was electrostatic. ATB-derived sequences were strongly shifted towards high net positive charge, with their net-charge distribution centred around +12 (interquartile range roughly +10 to +14), whereas database AMPs clustered near +3 (interquartile range ~+2 to +5). Consistent with this, ATB peptides exhibited markedly higher isoelectric points, with medians around pH 12 compared to ~pH 11 or below for database peptides.

This systematic bias towards highly cationic, high-pI sequences mirrors the enrichment in Lys and Arg observed at the sequence level and provides a clear electrostatic rationale for the robust activity of the synthesised ATB peptides against Gram-negative bacteria, whose outer surfaces are densely anionic.

Hydrophobicity- and amphiphilicity-related descriptors revealed a complementary but distinct pattern. On a normalized hydrophobicity scale, ATB peptides were on average more hydrophobic than database AMPs, with median values around 0.3 and few strongly hydrophilic sequences, whereas reference AMPs spanned a broader range from clearly hydrophilic to more hydrophobic compositions. In contrast, descriptors that capture the geometry of amphiphilicity rather than its global magnitude pointed to less classical helical segregation in the ATB set. The normalized hydrophobic moment and linear moment were both lower in ATB peptides than in database AMPs, and the angle subtended by hydrophobic residues was concentrated around small values (~30°) compared with much larger angles (typically ~60–150°) for many reference AMPs. Together, these metrics suggest that while ATB peptides are globally hydrophobic enough to interact with membranes, their hydrophobic residues are organised into tighter or more localised patches rather than the extended, large hydrophobic faces typical of canonical α-helical AMPs. At the same time, predicted membrane penetration depth was shifted towards larger values for ATB peptides (with medians at or near the upper end of the scale), indicating that these sequences are, in aggregate, expected to insert more deeply into the bilayer than many database AMPs, despite their more compact hydrophobic patterning.

Descriptors associated with conformational behaviour and self-association showed more nuanced differences. On the disorder-related scale, ATB peptides displayed a distribution shifted towards lower values compared with database AMPs, indicating a systematic change in predicted conformational propensity; assuming higher values on this scale correspond to greater intrinsic disorder, this would imply that ATB peptides are, on average, somewhat less disordered and more prone to adopt defined conformations than many known AMPs, although this interpretation depends on the exact calibration of the underlying metric. The propensity to *in vitro* aggregation was highly skewed for both groups, with medians close to zero but long upper tails. ATB peptides exhibited a somewhat higher upper quartile and mean than the database set, suggesting that while most ATB candidates are not strongly aggregation-prone, a subset could be more prone to self-association under physiological conditions—an aspect that may impact both potency and toxicity and will require experimental follow-up.

Although these physicochemical descriptors were computed for the broader set of APEX-predicted ATB peptides, the 24 synthesised molecules were deliberately selected from within this high-charge, moderately hydrophobic regime. Their sequences combine very high cationic content with varying degrees of hydrophobic and aromatic enrichment (for example, multiple Trp, Phe and Leu/Ile residues in ATB5, ATB6, ATB9, ATB20 and ATB60), making them representative of the dominant trends observed in the global ATB distribution. The strong and often low-micromolar MICs observed for most of these peptides against a panel of Gram-negative and Gram-positive pathogens are therefore consistent with the joint profile of extreme positive charge, elevated isoelectric point, sufficient global hydrophobicity and predicted deep membrane penetration revealed by the physicochemical features analysis. At the same time, the relatively modest hydrophobic moments and small hydrophobic-face angles indicate that many ATB peptides likely do not conform to the archetype of long, perfectly segregated amphipathic helices commonly found in AMP databases. Instead, they appear to exploit more compact or patchy amphiphilic motifs, which nevertheless support robust antimicrobial activity. Taken together, the alignment between sequence composition, physicochemical bias and experimental activity suggests that AllTheBacteria, when coupled to APEX, naturally enriches for a distinctive class of highly cationic, deeply inserting antimicrobial peptides that partially overlap with, but also extend beyond, the physicochemical space of canonical AMPs.

### Supplementary Note 4. Membrane-induced structural transitions of ATB peptides revealed by circular dichroism

\we examined their secondary-structure propensities by circular dichroism (CD) spectroscopy and quantified the resulting spectra with the BeStSel algorithm. By comparing peptide behaviour across four environments of increasing membrane mimicry (water, 50% methanol, 60% trifluoroethanol, and 10 mmol $L^{-1}$ Sodium dodecyl sulfate), we sought to capture not only intrinsic structural preferences but also the extent to which each peptide undergoes folding transitions upon encountering amphiphilic or hydrophobic interfaces.

Across the panel, the peptides were predominantly unstructured in aqueous solution, exhibiting minimal helical content and substantial β-antiparallel and disordered fractions. This behaviour is fully consistent with their primary sequences: most ATB peptides are highly enriched in Lys and Arg, with limited hydrophobic continuity and low predicted hydrophobic moments, features that disfavour stable secondary structure in water. Even among the peptides that later proved to be the strongest antimicrobial candidates, such as ATB3, ATB5, and ATB9, the CD signatures in water reflected largely disordered ensembles. Such conformational plasticity is common among potent host-defence peptides, many of which only adopt ordered structures upon binding to bacterial membranes.

Introduction of 50% methanol produced only modest structural stabilization, with small increases in α-helicity and slightly reduced β-antiparallel contributions. The limited response in this intermediate solvent underscores a central feature of the ATB sequences: although they contain hydrophobic residues, these are arranged in short or discontinuous patches rather than in the extended hydrophobic segments necessary for robust helix nucleation in partially structure-promoting environments. As a result, most ATB peptides remain largely disordered until they encounter a stronger membrane mimic.

By contrast, substantial helix formation emerged in the presence of TFE and SDS, particularly for peptides containing aromatic or aliphatic hydrophobic residues interspersed within their cationic frameworks. In SDS, a subset of peptides—including ATB3, ATB5, ATB6, ATB9, ATB20 and ATB60—displayed pronounced increases in α-helix content, in some cases exceeding 30–50% helicity. The same peptides exhibited reductions in β-antiparallel structure and a clear redistribution of their secondary-structure profiles toward canonical membrane-bound helical states. ATB5 was especially notable, undergoing a dramatic transition to ~50% helix in SDS, consistent with its mixed hydrophobic–aromatic composition and its high antimicrobial potency.

The peptides that remained largely unstructured even in SDS, such as ATB1 and ATB15, share strikingly simple sequence architectures dominated by short-period Lys/Arg repeats with minimal hydrophobic residues. These low-complexity sequences not only show weak helical inducibility but also demonstrate limited antimicrobial activity, supporting a direct link between sequence architecture, membrane-responsive folding, and biological function.

### Supplementary Note 5. Cytotoxicity assays show ATB peptides are generally not cytotoxic

To evaluate the safety profile of the ATB peptides, we measured their cytotoxicity against human embryonic kidney (HEK293T) cells using a 3-(4,5-dimethylthiazol-2-yl)-2,5-diphenyltetrazolium bromide (MTT) viability assay and determined half-maximal cytotoxic concentrations ($CC_{50}$). The resulting values spanned several orders of magnitude, from low micromolar for the most cytotoxic candidates to extremely high nominal $CC_{50}$ estimates for the least toxic peptides. Across the 24-member panel, the median $CC_{50}$ was ~30 μmol $L^{-1}$ (interquartile range ~10–230 μmol $L^{-1}$), reflecting generally moderate cytotoxicity with substantial heterogeneity between individual sequences.

A subset of peptides exhibited low micromolar $CC_{50}$ values, indicating a relatively higher propensity for mammalian cell toxicity. The most cytotoxic molecules in the panel included ATB25, ATB65 and ATB55 ($CC_{50}$ ≈ 5–6 μmol $L^{-1}$), followed by ATB13, ATB15, ATB5, ATB76, ATB18 and ATB60 ($CC_{50}$ ≈ 9–12 μmol $L^{-1}$). These peptides are strongly cationic but also relatively enriched in hydrophobic residues, especially in the case of ATB25, ATB55 and ATB65, which is consistent with an increased likelihood of interacting with and perturbing mammalian membranes. By contrast, a second group—including ATB12, ATB3, ATB39, ATB56, ATB6, ATB61 and ATB51—displayed intermediate $CC_{50}$ values (≈20–100 μmol $L^{-1}$), suggesting more favourable safety margins at concentrations close to their antimicrobial MICs. Finally, several peptides—ATB2, ATB9 and ATB20—were only weakly cytotoxic within the tested range, with $CC_{50}$ values on the order of hundreds of micromolar. For four peptides (ATB1, ATB63, ATB64 and ATB99), the estimated $CC_{50}$ values were exceptionally large (on the order of $10^{30}$–$10^{59}$ μmol $L^{-1}$), far above the rest of the panel, consistent with very low or undetectable cytotoxicity under the assay conditions.

To place these data in the context of antimicrobial efficacy, we calculated a simple selectivity index as the ratio $CC_{50}$ (HEK293T) / MIC against *A. baumannii* ATCC 19606. Among the peptides with measurable MICs against this strain, several showed favourable therapeutic windows, with selectivity indices ≥10. These included ATB20, ATB51, ATB9, ATB61, ATB12, ATB2, ATB56 and ATB5, with ATB20 and ATB51 standing out ($CC_{50}$/MIC ≈ 57 and 48, respectively), followed by ATB9 and ATB61 (≈30 and 29) and ATB12 and ATB2 (≈22 and 16). These molecules combine strong antibacterial activity with low to moderate HEK293T toxicity, marking them as particularly attractive leads for follow-up optimization. A second group—ATB3, ATB6 and ATB13—displayed intermediate selectivity (indices ≈4–7), suggesting that modest improvements in potency or reductions in mammalian toxicity could render them viable candidates.

In contrast, a set of peptides exhibited narrow or unfavourable selectivity windows. ATB25, ATB39, ATB55, ATB60, ATB65 and ATB76 all had $CC_{50}$/MIC ratios ≤3.8, with ATB60 and ATB76 falling below unity (≈0.38 and 0.64, respectively), indicating that concentrations needed to achieve full antibacterial activity approach or exceed those that compromise HEK293T viability. Notably, several of these peptides—ATB60, ATB76 and ATB65—also showed strong outer-membrane permeabilization in the NPN assay, suggesting that the same physicochemical features that promote efficient disruption of bacterial envelopes may also predispose them to off-target effects on mammalian membranes.

Supplementary Table 1. ATB peptides synthesized and validated experimentally.

| Peptide | Sequence |
|---|---|
| ATB1 | MRLRLRLRLRLRLRLRLRLRLRLRLRLRLRLRLRLRLR |
| ATB2 | MKKLKIKKLKIKKLKIKKLKIKKLKSGLRSQ |
| ATB3 | MSLRKKLFVKTMLRVKIKKQSKQRLQKTKRLLQKLKRPWM |
| ATB5 | MLKLLSIKLKSKKKILRMTLTKLNKWLKTKKKWLRNLKTM |
| ATB6 | MKLKKLIKFFAFAKGISLLSRRPVRKRTGGLLKKYLAWKYLRRR |
| ATB9 | MRRRGWKVRLKRRRGWKVRLMRRRGWKVRLKRRRLVLPVWA |
| ATB12 | MIKVSKLVVAKKLNLRKKLKKYLKKPKLGKILLL |
| ATB13 | MLKGLKKFKWLKKLKRRLMELNELKKFKKLMELRS |
| ATB15 | MKKLKIKKLKIKKLKIKKLKIKNLKIKKLKIKNLKI |
| ATB18 | MRSLKRQLLKHKAVKLSIKKWQFLRKLKLVPKRH |
| ATB20 | MKLWKIVRFLALSKGAKALRKKPVRKAAASFVKSYALWKYIRRK |
| ATB25 | MTKHAIAAKSLLKKWIISRPLNMKKKSSPFQKIRKMFKRFFK |
| ATB39 | MLGRKGLRSVWNGRLNVRKRRQNVRLRKNVRRKNVPKPSVWLI |
| ATB51 | MTKLKMYLKKVRKNANNSTKQWKYGLPVRNAKLWNNFQPKK |
| ATB55 | MSLKKIAKLLANKKVRKIAWMLISHPKFIKMLKNSIKRR |
| ATB56 | MKILKKNMHNKILPMVSHKKWMMLIKRLKTLGRIKK |
| ATB60 | MFKTNLSRVSVKRLLIWVVWWLRLKKLKKKAKKFRKK |
| ATB61 | MLKLLKKLQMHLLKKLQKEKLLLNLQKKTLKRKRFRHA |
| ATB63 | MNKIILRVKKMLKNIKLKNNGIRKYNVKMEVKLVYFK |
| ATB64 | MRKMKYVYYHLLSRLYKRQGKWQSEKFVRLINKRNMYK |
| ATB65 | MKKLIRNIIYSKLKFLWRWHRPLIIQRRNL |
| ATB76 | MRIKLHLRLAFLRFRKKKSLSRLKPISKIRSK |
| ATB94 | MTLKMRPLIKLFTRKKERKKQIFSLKYFKKFKLIKTTK |
| ATB99 | MIMATKLIKKNKFAKELFSRKYKPRVIKPKKGKGSFKRKKR |
